# A Minimally Invasive, Scalable and Reproducible Neonatal Rat Model of Severe Focal Brain Injury

**DOI:** 10.1101/2025.11.10.687751

**Authors:** Victor Mondal, Emily Ross-Munro, Gayathri K. Balasuriya, Ritu Kumari, Isabelle K. Shearer, Andjela Micic, Abdullah Al Mamun Sohag, Alan Shi, Mikaela Barresi, David R. Nisbet, Glenn F. King, Richard J. Williams, Pierre Gressens, Flora Y Wong, Jeanie L.Y. Cheong, David W. Walker, Mary Tolcos, Bobbi Fleiss

## Abstract

**Background:** Neonatal brain injuries such as stroke cause focal ischemic lesions that often result in lifelong neurological disabilities, as treatment options are limited. To speed up the discovery of potential therapies, early-phase screening with models that reliably reproduce brain injury, with scalable injury volume and minimal confounders, such as varying anaesthesia duration and painful procedures, is essential.

**Methods:** Postnatal day 10 Sprague–Dawley rats of both sexes, with four litters per group and timepoint, were randomly allocated to delivery of intraperitoneal Rose Bengal (25, 40, or 60 mg/kg) and 10 minutes of light-emitting diode illumination through the intact scalp and skull. Infarct progression and reproducibility were assessed at 24 hours, 7 days, and 14 days post-injury. Outcomes included infarct volume and sensorimotor function, and cleaved caspase-3, glial fibrillary acidic protein (GFAP), and ionised calcium-binding adaptor molecule 1 (Iba1) immunoreactivity, with analysis of sex differences. Data were analysed using one-way or two-way ANOVA with Sidak’s post-hoc tests.

**Results:** There was no mortality due to the infarct, and procedure time was approximately 19 minutes across all groups; the lesion was consistent and supported scalability. The 25 mg/kg dose produced a reproducible cortical infarct (3.74 ± 0.58 mm³; CV = 31%). Lesion size increased with dose and decreased over time (11.15 ± 0.63 mm³ at 60 mg/kg versus 0.05 ± 0.007 mm³ at 14 days; p < 0.0001). Cleaved caspase-3 and glial activation persisted for 14 days, indicating ongoing apoptosis and gliosis. No sex-dependent effects were observed in lesion volume, behaviour, or gliosis.

**Conclusions:** This refined neonatal photothrombotic ischaemia model is reproducible, scalable, and ethically improved, requiring no skin incision. Its minimal surgical burden, absence of mortality, consistent histopathology, and measurable functional outcomes make it an ideal platform for preclinical screening of neuroprotective and reparative interventions in the developing brain.

## Introduction

Injury to the developing brain is the most common cause of permanent disability worldwide ^1^. Infants with the most severe focal brain injury, particularly those linked to focal ischemia, are at high risk of significant lifelong disability. For instance, most infants who suffer a stroke in the region of the middle cerebral artery in the first 28 days of their postnatal lives develop cerebral palsy, and over 80% will exhibit language, hearing, or visual field impairments ^2,3^. In addition, following severe hypoxic-ischemic encephalopathy (HIE), 25% of infants develop cerebral palsy, and 70% of those without a motor deficit will experience significant academic challenges ^4^. Among children under one year of age, accidental or abusive head trauma is the leading cause of traumatic brain injury (TBI)-related death and hospitalisation ^5,6^. These TBIs are often associated with large, focal lesions on clinical imaging ^7^, and more than 70% of infants with severe TBI who survive live with adverse cognitive and motor outcomes ^8^.

Traditionally, HIE and TBI are considered diffuse injuries, but in their most severe presentations, both involve focal cortical or subcortical infarctions, which can be caused by ischemia ^9^. Currently, there are no curative therapies for severe focal brain injuries. Therapeutic hypothermia, the current standard of care for HIE, is only partially effective, has limited applicability to other types of brain injuries ^10,11^ and is effective only in a resource-intensive setting ^12–14^. Severe brain injury is dramatically more prevalent in regions where there are inequalities in access to health care ^12^. This highlights the need for preclinical models that reproduce focal cortical lesions in the neocortex to support the development of therapies, including first-line screening models. Addressing this need requires models that are reproducible, ethically refined, and scalable across different injury severities, especially for early-phase therapy screening^15^.

Whilst several models exist to study focal brain injury, these have limitations for early-phase screening, especially when fixed scalability is required. The middle cerebral artery occlusion (MCAO) model is widely used for its clinical relevance, but infarct size varies widely across animals due to differences in collateral blood flow^16^, and mortality can reach 40%^17,18^. This limits reliable scalability and demands large group sizes. The endothelin-1 model has lower mortality but still shows inconsistent lesion size ^19^. The Rice-Vannucci model, commonly used for HIE, is similarly limited by variable lesion territories despite lower mortality (5–10%) ^20^. All three models are invasive, technically complex, and poorly suited for high-throughput early-phase neurotherapy screening.

In contrast, the photothrombosis model has a low mortality rate (<2%) and a highly reproducible lesion territory ^21–23^. Rose Bengal, a visible-light photoredox catalyst, is administered into the bloodstream, followed by targeted light exposure onto the brain to induce platelet aggregation within cerebral blood vessels, resulting in focal vascular occlusion. In the adult, the scalp, skin, periosteum, and skull cause scattering and attenuation of light that necessitate at least an incision to apply the light directly to the skull ^21^, or on the brain following a craniotomy ^23^. The first iteration of this model in the neonatal rat required an injection of Rose Bengal into the exposed jugular vein and the application of light directly to the skull over the cortex after a skin incision ^24^. Since that time, there have been substantial improvements to reduce invasiveness, including intraperitoneal injection of Rose Bengal and only a skin incision for light placement ^25^.

To the best of our knowledge, no minimally invasive protocol (Rose Bengal, intraperitoneal, no skin incision) with a fixed procedure duration has been established that correlates the Rose Bengal dose with highly reproducible infarct size and subsequent structural and functional damage. This is a critical gap, particularly for early-phase therapy development, where reproducibility and ethical refinement are essential, and for testing therapies such as hydrogel-based delivery systems and tissue transplants, where treatments are tailored to specific lesion characteristics ^26,27^. Moreover, standardising surgical duration is important for comparing injury severity, especially as anaesthesia itself increases cell death in the neonatal brain ^28^.

Here, we established a minimally invasive, fixed-duration, scalable and highly reproducible method for inducing photothrombotic focal brain injury in neonatal rats, and characterised the temporal progression of glial activation, cell death, and associated behavioural deficits. We hypothesised that higher doses of Rose Bengal would lead to more severe structural and functional outcomes. We examined sex differences in injury outcomes, given that male infants are known to have worse neurological outcomes than females ^29^.

## Materials and methods

### Animals and surgical procedures

Please also refer to the **Supplementary Methods** for further information. All experiments were approved by the RMIT University Animal Ethics Committee (AEC 1918) and are reported in accordance with the ARRIVE guidelines. Pregnant Wistar rats were obtained from Australian Bioresources (Perth, Western Australia) and allowed to deliver undisturbed. Pups were weighed on postnatal day (P) 8 and randomly assigned to treatment groups within litters using a restricted block design. P10 was chosen for the induction of focal cortical injury as it is developmentally equivalent to a term-born human infant ^30^.

Predetermined exclusion criteria included humane endpoints and technical artefacts; only one pup (sham group) died before its endpoint. For histology, one pup per litter per timepoint was analysed; for behavioural assays, each pup was treated as an independent replicate. Sex differences were assessed by adding additional animals from independent litters to the 25 mg/kg group.

P10 rats were anaesthetised with 5% isoflurane in 0.4 L/min O_₂_ (2–3% maintenance) and kept normothermic on a thermistor-controlled pad (Thorlabs, USA). Rose Bengal (Merck, Australia; cat#330000) was injected intraperitoneally at 25, 40, or 60 mg/kg, based on a literature review ^23^. After 5 minutes, a 1 mm LED light source (Thorlabs M565F3, 565 nm) was positioned 2 mm right of Bregma and illuminated through the intact scalp and skull for 10 minutes (**Supplementary Figure 1**). The full procedure averaged 19 minutes for all groups.

At completion, buprenorphine (0.05 mg/kg) and meloxicam (5 mg/kg) were administered subcutaneously. Pups recovered for 1 hour on a heating pad before being returned to the dam. Sham controls received Rose Bengal without illumination to control for dye effects ^31^. Light exposure alone produces no detectable lesion and was not considered a confound. Further procedural details and anatomical schematics appear in *Supplementary Methods* and *Supplementary Figure 1*.

### Behavioural testing

Behavioural testing was conducted between 0800 and 1200 hours by blinded assessors. Testing epochs corresponded to: P11 (24 hours post-injury) – acute neuroinflammatory phase; P17 (7 days) – subacute consolidation/remodelling; P22–P24 (12–14 days) – early recovery, coinciding with rapid myelination and circuit maturation, approximating 2–3 years human age ^30,32,33^.

#### Wire hang test

At P11 and P17, rats grasped a wire using both forepaws and were allowed to hang for ≤5 minutes. Three trials (30 seconds apart) were video-recorded, and the total hang time was summed.

#### Cylinder rearing test

At P22 (12 days post-injury), rats were filmed for 5 minutes in a transparent cylinder (185 mm height × 130 mm diameter). Forepaw contacts were classified as contralateral, ipsilateral, or bilateral, and contralateral use (%) = (contralateral touches / total touches) × 100.

#### Adhesive-tape removal test

At P22, adhesive discs (0.95 cm diameter; Merck Z743588) were applied to both forepaws. Each rat completed 3 trials (maximum 3 minutes per trial, 1-minute interval). The time to contact and remove each tape was recorded.

#### Open-field test

At P11 and P17, rats explored a 30 × 30 cm arena for 3 minutes under overhead video. Parameters included locomotor activity, centre time, rearing, and immobility.

All behavioural paradigms are further detailed in **Supplementary Methods** and schematised in **Supplementary Figure 2**.

### Brain tissue collection and processing

Brains were collected at P11, P17, and P24 (**Supplementary Figure 2**). Animals were anaesthetised with Lethabarb (>200 mg/kg, Virbac Australia) + lignocaine (10 mg/mL) and perfused with PBS followed by 4% paraformaldehyde (PFA). Brains were post-fixed for 12–18 hours, transferred to 70% ethanol, and processed (Leica ASP300S) into paraffin. Coronal sections (8 µm) were cut from Bregma +3.24mm to beyond the lesion. Three sections per slide were mounted on Superfrost Plus slides (each section spaced 160 µm apart) and stored for analysis.

### Histological analyses

#### Haematoxylin and eosin (H&E)

Every 20th section was stained to define lesion morphology. Area and depth were measured in ImageJ as described previously ^34,35^. Infarct volume was estimated using Cavalieri’s principle (V = ΣA × P × T). Lesion location was referenced to the Paxinos and Watson atlas ^36^.

#### Reproducibility and cortical atrophy

Lesion consistency between litters was quantified as the coefficient of variation (CV) ^37^. Cortical atrophy = [1 – (ipsilateral/contralateral area)] × 100, averaged across sections, as previously ^35,38^. Ventricular enlargement was expressed as the ipsilateral-to-contralateral (LVR) ratio.

### Immunohistochemistry and immunofluorescence

IHC and IF were performed as previously described ^39,40^, with full details in **Supplementary Methods.** Sections were dewaxed, subjected to citrate buffer antigen retrieval (pH 6.0), and blocked in 5% normal goat serum with 0.2% Triton X-100. Primary antibodies: anti-GFAP (1:1000, Dako Z033401-2), anti-Iba1 (1:1000, Novachem 019-9741), anti-cleaved caspase-3 (1:500, Cell Signalling 966L1). Secondary labelling used either donkey anti-rabbit Alexa Fluor 488 (1:500, Invitrogen, Cat # A32790), biotinylated anti-rabbit, goat polyclonal (Vector Laboratories, Cat# VEBA1000), followed by biotin-streptavidin ABC with DAB visualisation. Slides were counterstained with DAPI or mounted in DPX as appropriate.

### Image acquisition and quantification

All analyses were performed blinded to group. Slides were imaged using an Olympus VS120 scanner (10x for H&E; 20x for IHC/IF). For each marker, three sections per brain were analysed at the lesion maximum and ± ⅓ lesion span from either side. Regions of interest included the infarct core, border, and corresponding sham cortex (**Supplementary Figure 3**). As previously ^34,38,41^, positive area (GFAP, Iba1) was quantified via pixel thresholding after background subtraction; CC3-positive cells were manually counted in matched ROIs with full details in ***Supplementary Methods***.

### Statistical analyses

Data are presented as mean ± SEM. Analyses were performed in GraphPad Prism (v8, USA). A *p* < 0.05 was considered significant. One-way and two-way ANOVAs with Sidak’s post hoc tests were applied to behavioural and histological data, with the specific models described in the figure legends. Lesion metrics and atrophy were tested by two-way ANOVA; correlations between lesion volume and behaviour were assessed by Pearson’s *r*. Sex effects were analysed using two-way ANOVA with Sidak’s correction. Power analyses used G*Power (v3.1) ^42^. Full statistical outputs, inclusion/exclusion criteria, and sample size rationale are reported in **Supplementary Methods** and **Supplementary Tables**.

## Results

### Severe focal cortical injury did not affect body weight or cause mortality

Across all litters (n = 4 per timepoint), body weight increased steadily from P10 to P22 in both sham- and Rose Bengal–treated pups (**Supplementary Table 1**; **Supplementary Figure 4**). Sham animals gained from 19.6 ± 0.6 g at P10 to 53.3 ± 2.3 g at P22, and comparable growth trajectories were observed in the 25, 40, and 60 mg/kg dose groups. Only 1 of 54 pups in this study died: a sham rat, 24 hours after surgery.

### The severity of the injury was proportional to the Rose Bengal dose

#### Infarct volume is maximal at 60 mg/kg

A two-way ANOVA revealed significant effects of Rose Bengal dose and time after injury across pups (*n* = 36; 4 pups/group per timepoint; **Figure 1A, B**; **Supplementary Table 1,3**). The infarct volume was largest at 24 hours and smallest at 14 days post-injury for all Rose Bengal doses, with higher Rose Bengal doses producing larger infarcts (**Figure 1A-B).**

**Figure 1.**
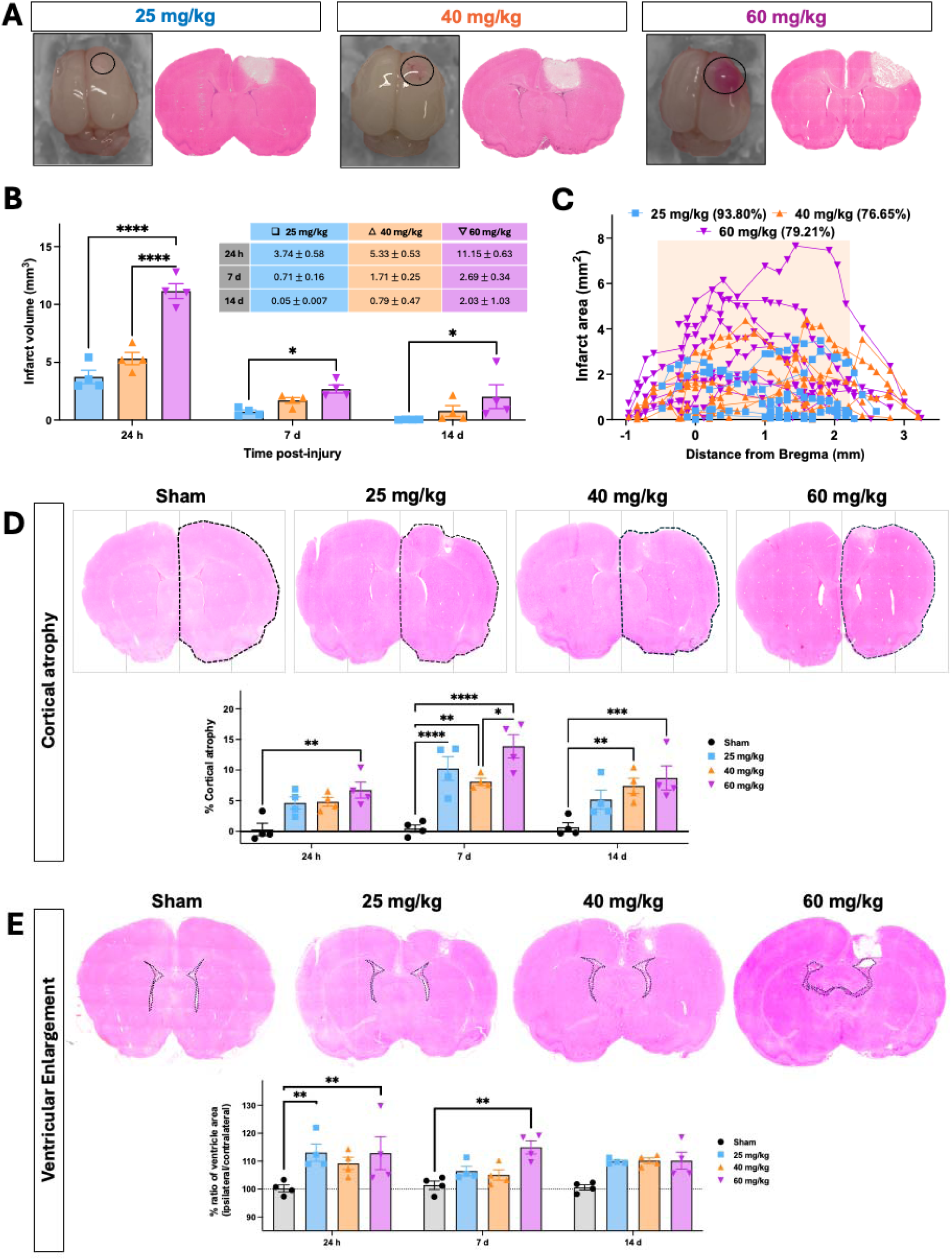
Dose-dependent relationship of Rose Bengal with infarct volume, cortical architecture, and secondary injury. (**A**) Macroscopic images of Rose Bengal-induced lesions (left) and representative H&E-stained coronal sections (right) showing dose-dependent infarct size and depth across 25, 40, and 60 mg/kg groups. (**B**) Quantification of infarct volumes at 24 hours (h), 7 days (d), and 14 days across Rose Bengal doses (mean ± SEM; n = 4 per timepoint per group). Two-way ANOVA with Sidak’s post-hoc test was used for multiple comparisons. *p < 0.05, ****p < 0.0001. Inset = lesion volume in millimetres cubed; ± plus or minus standard error of the mean. (**C**) Infarct area plotted along the anterior-posterior axis relative to Bregma. Orange shading denotes the region of the primary motor cortex (+2.2 mm to −0.5 mm from Bregma). Each line represents one animal (*n* = 12 per dose from four litters; see Supplementary Table 1). (**D–E**) Representative H&E-stained sections from sham, 25, 40, and 60 mg/kg groups, highlighting (**D**) cortical atrophy and (**E**) ventricular enlargement and quantification of (**D**) cortical atrophy and (**E**) ipsilateral ventricular volume across time points and doses (mean ± SEM; *n* = 4 pups per group from four litters). Two-way ANOVA with Sidak’s post-hoc test was used for multiple comparisons. *p < 0.05, **p < 0.01, ***p < 0.001, ****p < 0.0001.

#### The infarct is highly reproducible and consistently localised to the motor cortex in the 25 mg/kg Rose Bengal group

In the 25 mg/kg injury group, 93.8% of the infarct was in the primary motor cortex (**Figure 1C**). At 40 mg/kg and 60 mg/kg, the percentage within the motor cortex was 76.6% and 79.2%, respectively. Lesions were induced in all animals treated with Rose Bengal. Lesion reproducibility between litters was assessed at 24 hours after injury by calculating the CV for the lesion volume (CV_lesion_), and the lesion area as a percentage of the hemisphere (CV_percent_: **Supplementary Table 2**). The CV was the lowest (i.e., the least variation between litters) in the 60 mg/kg group (CV_lesion_, 11% - CV_percent_ 16%), although, as outlined above, 30% of the lesion extended beyond the motor cortex (**Figure 1C**). CV values across all groups ranged from 11 to 31%, within the previously defined high-reproducibility range ^43^ (**Supplementary Table 2**).

#### Infarct area correlates positively with infarct depth

A Spearman correlation coefficient test revealed a significant association between infarct area and depth at all Rose Bengal concentrations (**Supplementary Figure 5**): 25 mg/kg Rose Bengal dose (r_s_ = 0.729; p < 0.0001), 40 mg/kg (r_s_ = 0.704; p < 0.0001) and 60 mg/kg (r_s_ = 0.669; p < 0.0001). Although Rose Bengal injury was primary cortical, subcortical white matter damage was observed in the 40 mg/kg and 60 mg/kg groups 24 hours after injury.

#### Cortical atrophy and ventricular dilation were dose-dependent

There was a significant effect of Rose Bengal dose and time after injury on cortical atrophy in the two-way ANOVA (**Figure 1D, Supplementary Table 4**). Cortical atrophy significantly increased at day 7 and 14 post-injury compared to sham values (**Figure 1D).** There was also a significant effect of Rose Bengal dose on ventricular volume in the two-way ANOVA (**Figure 1E Supplementary Table 4**). Specifically, the total volume of the ipsilateral ventricles was significantly larger at 24 hours and 7 days post-stroke compared to sham-operated controls in the multiple comparison testing (**Figure 1E**).

### Severe focal injury in the neonate caused acute cell death and gliosis

At 24 hours post-injury, the number of CC3-positive cells and area coverage of Iba1 and GFAP were quantified within the infarct border area. We observed a significant increase in CC3 cell density (**Figure 2A**), Iba1 area coverage (**Figure 2B**) and GFAP area coverage (**Figure 2C**) at the infarct border in all injury groups compared with sham; statistical outputs in **Supplementary Table 5,6**. There were no significant differences between the 25, 40, and 60 mg/kg groups in the width of the glial reactive zone for Iba1-positive cells (one-way ANOVA, **Supplementary Figure 6**).

**Figure 2.**
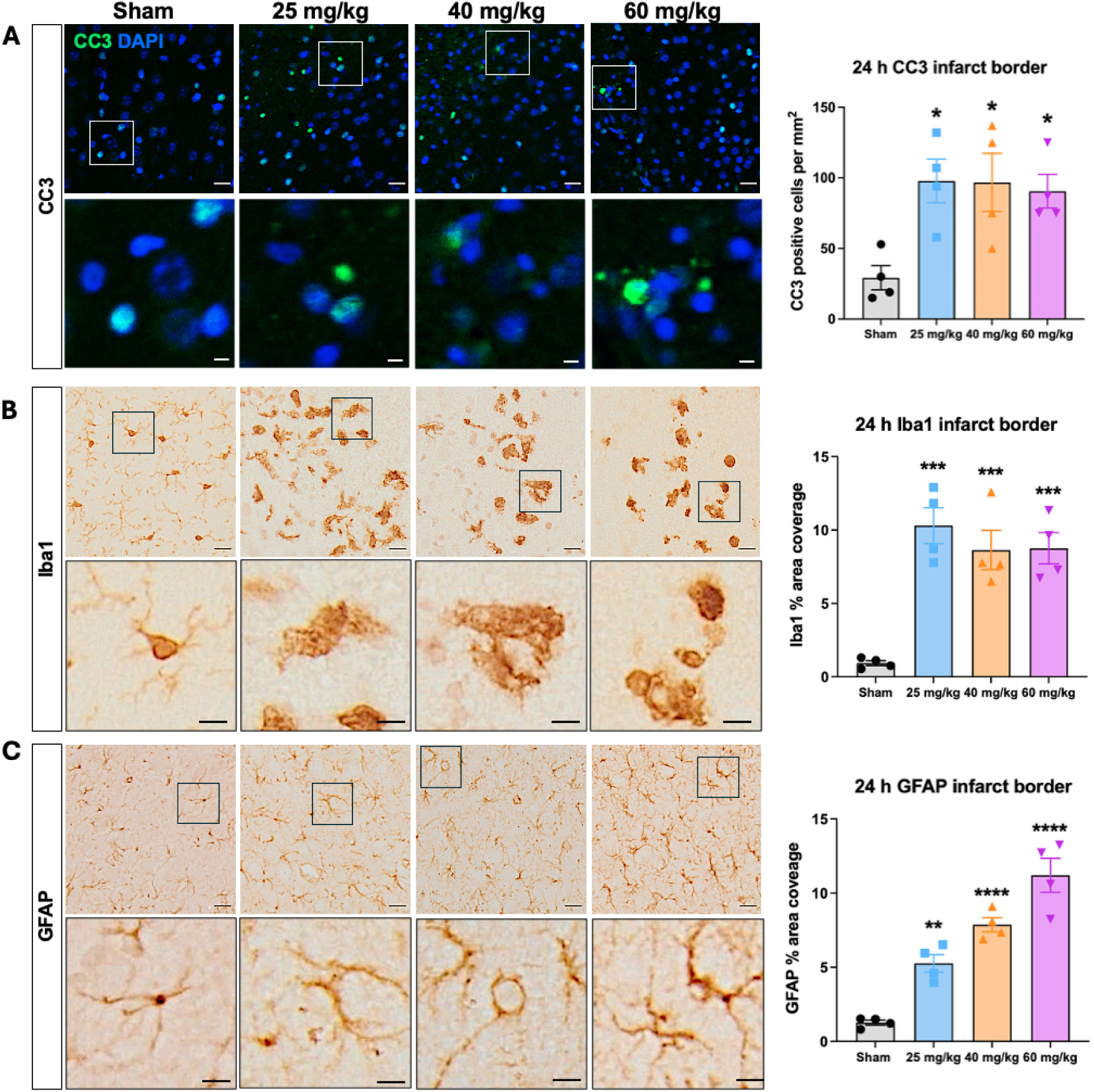
Cell death and glial reactivity in the infarct border zone. Representative images of (**A**) CC3, (**B**) Iba1 and (**C**) GFAP immunostaining in the infarct border at 24 hours (h) post-injury and sham surgery, paired with quantification of total cell number per mm^2^ or the percent of glial reactive area covered. Data are presented as mean ± SEM (n=4 litters; n=4 pups per group). One-way ANOVA with Sidak’s *post-hoc* test was used for multiple comparisons. Scale bars, upper = 20 µm, lower = 5 µm; *p < 0.05, **p < 0.01, ***p < 0.001, ****p < 0.0001.

### Cell death and gliosis persisted after severe focal injury in the neonate

CC3, Iba1 and GFAP immunoreactivity were significantly increased in the infarct core at 7-and 14-days post-injury across all injury doses compared to sham controls based on a two-way ANOVA (**Figure 3A-C; Supplementary Tables 5,7**). CC3, Iba1- nor GFAP-positive cells in the core showed significant changes 24 hours after injury. Microglia/macrophage and astrocytes are recruited into the infarct core at later time points (7- and 14-day post-injury) following early localisation to the glial reactive zone at the border of the lesion.

**Figure 3.**
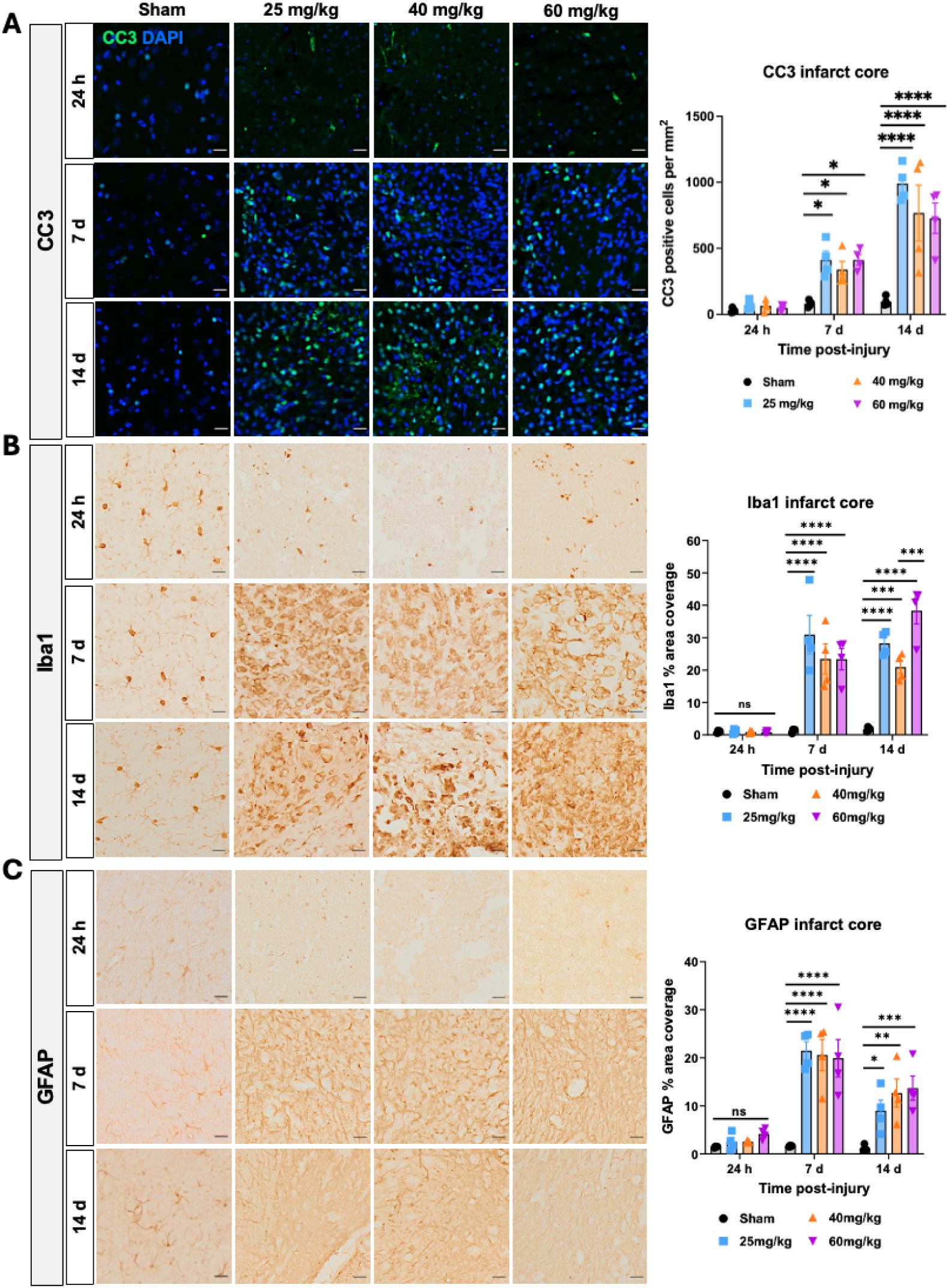
Cell death and glial reactivity in the infarct core. Representative images of (**A**) CC3 (**A**) Iba1 (**B**) and GFAP (**C**) in the core at 24 hours (h), 7 days (d) and 14 days after injury together with quantification of the cell density of CC3 and percent of infarct core area covered by Iba1 and GFAP. Data presented as mean ± SEM. *n* = 4 litters: *n* = 4 pups per group. Analysis using two-way ANOVA followed by Sidak’s *post-hoc* test for multiple comparisons. *p < 0.05, **p < 0.01, ***p < 0.001, ****p < 0.0001. Scale bars = 20 µm.

### Severe focal cortical injury in neonate rats impaired wire hang time and contralateral forepaw use in the days and weeks after injury

#### Wire-hang test

Injury significantly affected total hang time at 24 hours and 7 days (one-way ANOVA). Sham pups hung longer than the 25 mg/kg group at 24 hours (15.7 ± 2.2 seconds versus 8.6 ± 0.9 seconds; p = 0.039). There was also a trend toward reduced hang time at 24 hours for the 40 mg/kg group (p = 0.058) and at 7 days for the 25 mg/kg group (p = 0.069; **Figure 4B–C; Supplementary Table 8**).

**Figure 4.**
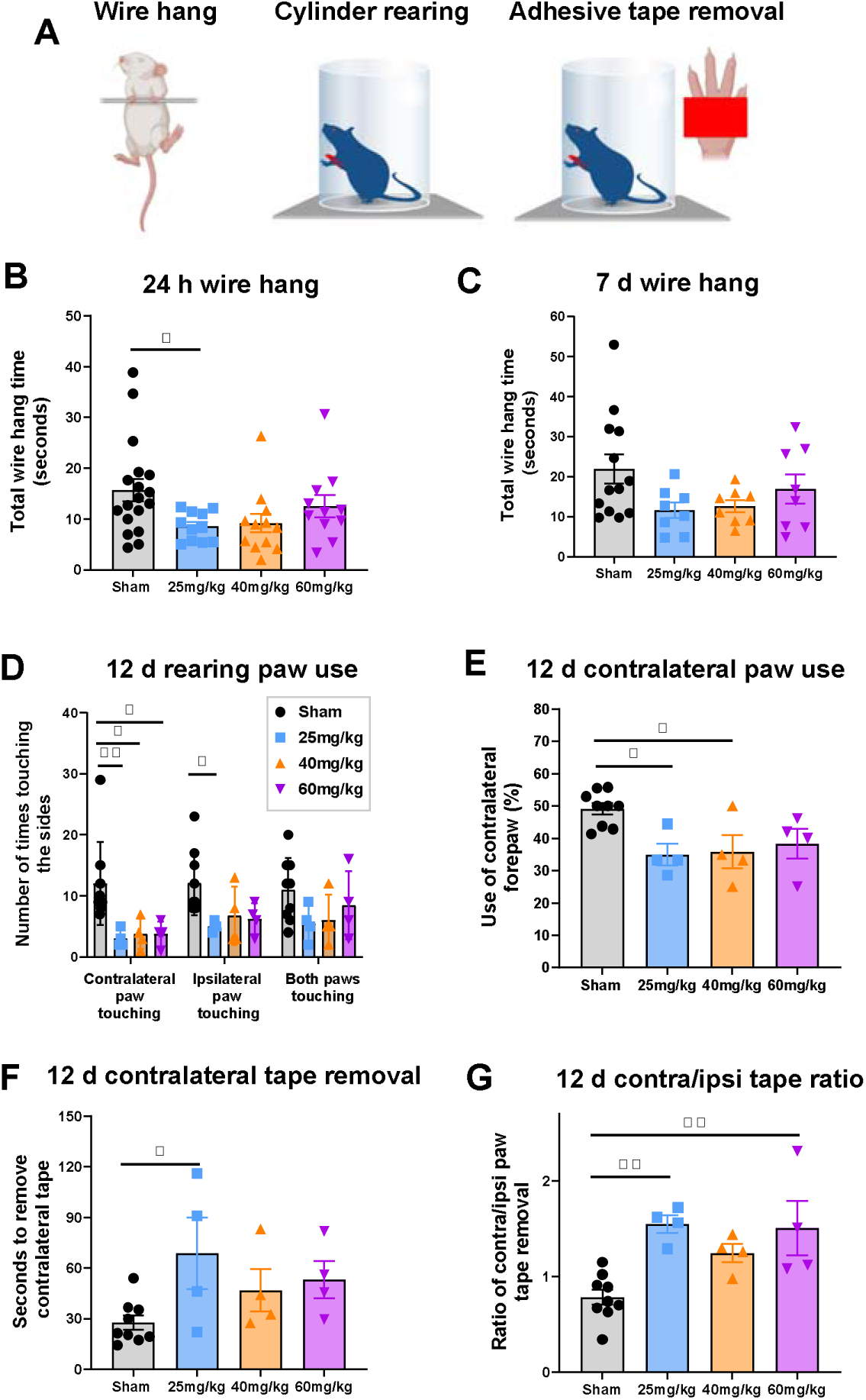
Severe focal injury led to persisting behavioural deficits. Changes were assessed using the (**A**) wire-hanging test, cylinder rearing and adhesive tape removal tests. Wire hang data are shown at (**B**) 24 hours (h) and (**C**) 7 days (d) post-injury. Cylinder rearing data include (**D)** use of contralateral, ipsilateral, and both forepaws and (**E**) percentage of contralateral forepaw use at 12 days after injury at postnatal day (P)22. Adhesive tape removal data show (**F**) time to remove tape from the contralateral forepaw and (**G**) the ratio of contralateral/ipsilateral forepaw use at P22. Data are presented as mean ± SEM. One-way and two-way ANOVA with Sidak’s post-hoc test was used for multiple comparisons *p < 0.05, **p < 0.01.

#### Cylinder rearing

Injury altered forelimb use (one- and two-way ANOVA). Pups that received 25 mg/kg of Rose Bengal showed reduced use of the contralateral and ipsilateral forepaw 12 days after injury, while those given 25 or 40 mg/kg displayed contralateral forepaw hypokinesia, with significantly lower contralateral use than sham (**Figure 4D–E; Supplementary Table 8**).

#### Adhesive-tape removal

Sensorimotor performance was impaired 12 days after injury (one-way ANOVA). Sham pups removed the tape faster (27.6 ± 4.2 seconds) than the 25 mg/kg group (68.7 ± 21.3 seconds; p < 0.05). No significant differences were found for the 40 mg/kg (46.8 ± 12.5 seconds) or 60 mg/kg (53.1 ± 11.0 seconds) groups (**Figure 4F**). The contralateral/ipsilateral tape removal ratio was increased in the 25 mg/kg (1.5 ± 0.1) and 60 mg/kg (1.5 ± 0.3) groups compared with sham (0.8 ± 0.1; p < 0.01), but not at 40 mg/kg (1.2 ± 0.1; **Figure 4G; Supplementary Table 8**).

No significant injury effects were observed in *open-field behaviour* (locomotion or time spent in the centre; **Supplementary Figure 7**).

### Exploratory correlation between lesion volume and motor function

We next explored whether lesion volume correlated with motor performance to identify the optimal Rose Bengal dose (**Supplementary Table 9**). Among the eight behavioural measures in Figure 4, six were significantly altered in the 25 mg/kg group but only two in the higher-dose groups, prompting a Pearson correlation analysis.

At 24 h, wire-hang performance showed a moderate, non-significant negative association with lesion volume in the 25 mg/kg (r = –0.53, p = 0.24) and 40 mg/kg groups (r = –0.72, p = 0.25), and no correlation at 60 mg/kg (r = –0.03, p = 0.49). Pooling the 25 and 40 mg/kg data increased power, revealing a strong trend between greater lesion volume and shorter hang time (r = –0.63, p = 0.06).

When wire-hang scores at 24 h were compared with lesion volume at 7 days, negative correlations emerged at 25 mg/kg (r = –0.81, p = 0.09) and 40 mg/kg (r = –0.99, p = 0.002), suggesting early motor deficits predict later tissue loss.

For long-term outcomes, adhesive-tape removal performance at 12 days correlated strongly and significantly with lesion volume at 14 days in the 25 mg/kg group (r = –0.95, p = 0.03), whereas only weak, non-significant relationships were found at 40 mg/kg (r = –0.17, p = 0.42) and 60 mg/kg (r = –0.16, p = 0.42).

Together, these findings indicate that moderate-dose photothrombosis (25–40 mg/kg) yields behaviour–lesion relationships consistent with scalable, reproducible injury suitable for functional outcome studies.

### The impacts of severe focal brain injury were not sex specific

To address whether sex influenced lesion characteristics or behaviour, additional animals were added to the 25 mg/kg group to give n of 7-14 (**Figure 5**). No significant differences were found between males and females in infarct volume (**Figure 5A**), wire hang at 24 hours or 7 days after injury (**Figure 5B**), anxiety-related behaviours (**Supplementary Figure 8**), CC3 positive cells (**Figure 5C)** or area coverage of Iba1 (**Figure 5D**) or GFAP (**Figure 5E**). Two-way ANOVA results are summarised in **Supplementary Table 10**. Power calculations were undertaken (G*Power, 3.1), **Supplementary Table 11**, to determine Cohen’s effect size and the number of rats required to detect statistically significant sex differences in a two-tailed t-test (α = 0.05, power = 0.8). For infarct volume, the number of pups required at 24 hours would be 29 pups, at 7 days would be 385, and at 14 days would be 80. For a wire hang at 24 hours to detect a difference between the shams would require 297 pups, and between stroke would require 232 pups; at 7 days, it would require 801 to detect a difference between the sham and 174 between stroke pups. Accordingly, it would not be practical or ethical to conduct additional studies to examine sex-dependent effects on these outcome markers.

**Figure 5.**
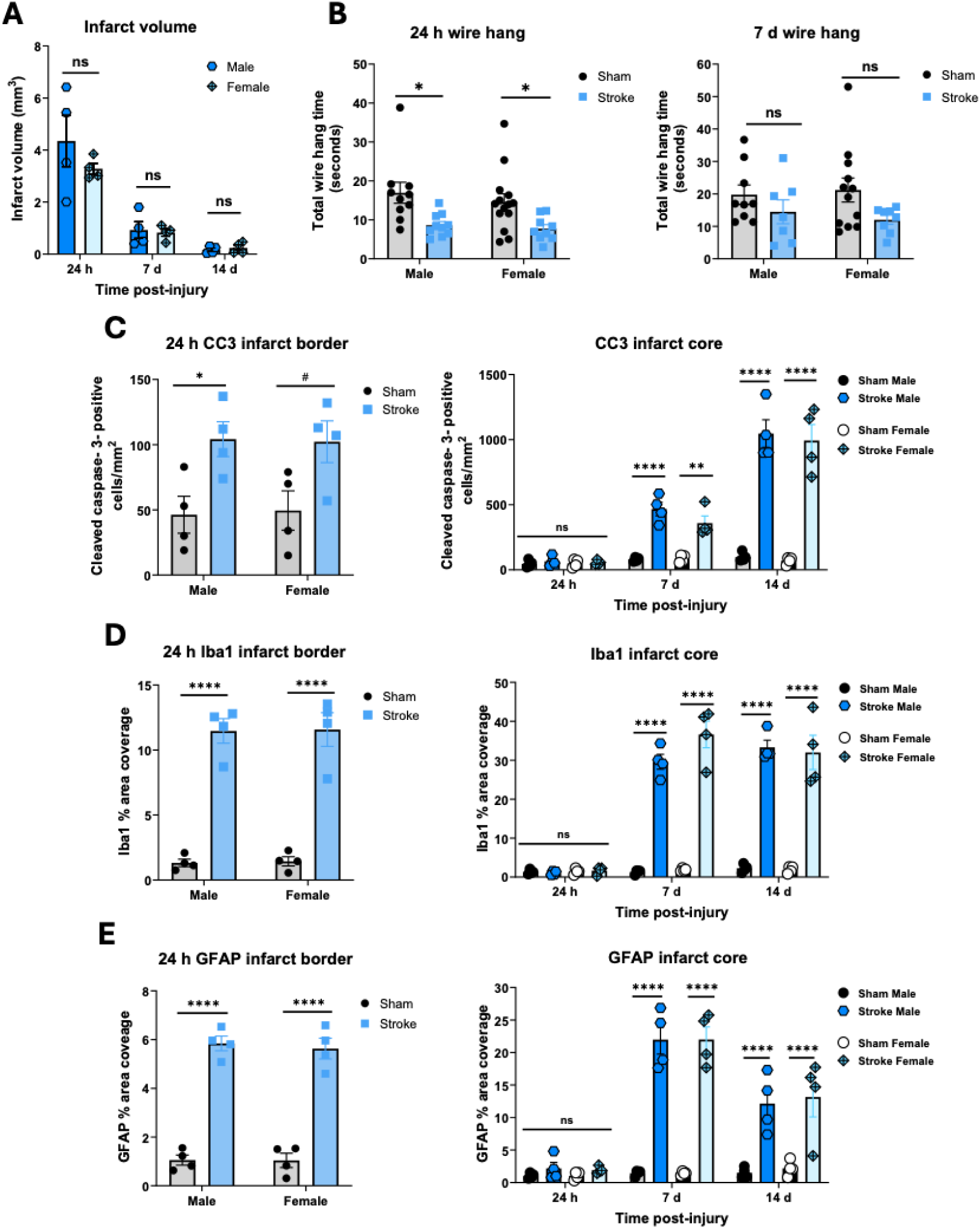
No sex differences in the effects of injury on behaviour and neuropathology. (**A**) Infarct volume quantified from H&E-stained brain sections at 24 hours (h), 7 days (d), and 14 days post-injury, (*n* = 4 per group). (**B**) Wire-hanging times at 24 hours post-injury for sham and injured males (*n* = 10), sham females (*n* = 14), and injured females (*n* = 9), and at 7 days post-injury in for sham males (*n* = 9), sham females (*n* = 12), injured males (*n* = 7), and injured females (*n* = 8). (**C**) CC3 positive cells in the infarct border at 24 hours and in the infarct core at 24 hours, 7 days and 14 days (*n* = 4 per group). Area coverage in the infarct border at 24 hours and in the infarct core at 24 hours, 7 days and 14 days for (**D**) Iba1 and (**E**) GFAP (all *n* = 4). Analyses were performed using two-way ANOVA with Sidak’s *post-hoc* test for multiple comparisons. *p < 0.05, **p < 0.01, ****p < 0.0001, ^#^p = 0.05-0.099; ns = not significant.

## Discussion

This study describes a minimally invasive, highly reproducible, scalable and specific form of motor cortex infarction in the neonatal brain that correlates with long-term functional impairment. A 19-minute surgery using a 25, 40 or 60 mg/kg dose of Rose Bengal activated by a 1 mm diameter light applied to the intact skin for 10 minutes resulted in a cortical lesion. The 25 mg/kg dose of Rose Bengal most reliably induced a lesion localised to the motor cortex and functional impairments, which correlated with lesion volume, a key factor in developing an early-phase screening model with translational potential.

Body weight did not differ between the sham and injury groups, likely due to the short procedure duration, the use of short-acting anaesthesia, and the modest lesion volume compared to other models. In the present study, lesion volumes at 24 hours post-injury across Rose Bengal doses ranged from 3.7 to 11.1 mm^3^. Knezic and colleagues reported larger infarct volumes, 8.2 to 14.0 mm^3^ that were associated with weight loss in young adult mice following photothrombotic stroke ^21^. Similarly, infarct size strongly correlated with body weight loss in an adult mouse MCAO model ^44^. The lack of systemic effects on weight is advantageous in a high-throughput screening model, since it avoids confounding effects associated with weight loss or general poor health. Weight loss is also common in MCAO models, where anaesthesia typically lasts ∼90 minutes ^45^ or repeated exposure is required^46^. Anaesthesia itself can cause weight loss, even without other interventions ^47^, and it can also lead to brain cell death ^28^. Neonates may tolerate moderate cerebral infarction without affecting their weight, since maternal care mitigates stress and provides better nutrition than in adults, who must obtain food and water independently ^48^.

An additional benefit of this model is that it avoids skin incision, thereby reducing overall harm to the animals, including post-operative burden, such as local inflammation and stress confounders. This contrasts with invasive neonatal photothrombotic models, which require jugular vein injection, midline scalp incision, and craniotomy ^24^ or, at a minimum, a skin incision to place the light source against the skull ^25^. In our protocol, light placement through the skin is possible because clear anatomical landmarks on the skull are visible, allowing reliable identification of Bregma. This minimally invasive approach also facilitates testing of therapies that themselves require surgical delivery at a different time or day, such as hydrogel implants^26,27^. Furthermore, there was no mortality in the infarct group, compared to a rate of 2.45% in a model requiring a skin lesion for light placement ^25^ and substantially lower than in models of MCAO, which can reach 40%^17^. Finally, this procedure is amenable to high-throughput screening; up to 20 pups can be processed per morning by two researchers, allowing for multiple therapeutic agents to be screened relatively quickly.

To assess biological reproducibility, we calculated the coefficient of variation (CV) for each litter, treating it as a biological replicate. This approach avoids pseudoreplication and captures variability across independent applications of the model, which is important given that up to 60% of the variance in brain injury models arises from between-litter effects ^49^. CVs are rarely reported in perinatal injury studies, but for the Rice–Vannucci model, a CV of 54% has been noted ^50^. In adult stroke models, a review reported CV ranges of 50–150% for intraluminal MCAO, 80–100% for embolic stroke, and 5–30% for photothrombotic models ^43^. The CV is a practical index of model utility in preclinical screening: to detect a 30% difference between groups (80% power, α = 0.05), 16 animals per group are required at a CV of 30%, but 44 are required at a CV of 50%. Importantly, smaller lesion volumes inflate CV, as even minor deviations significantly affect the ratio; conversely, larger lesions reduce CV because proportional variability is lower. Accordingly, the 60 mg/kg dose was expected to produce a lower CV, but even at 25 mg/kg, the CV was within the highly reproducible range.

The photothrombotic lesion model produced hemispheric asymmetry and ventricular enlargement, reflecting tissue loss and chronic atrophy, consistent with outcomes from other neonatal injury models ^24,51–53^. Higher Rose Bengal doses led to greater ipsilateral hemisphere volume loss and enlargement of the lateral ventricle. Importantly, in babies and children following neonatal stroke, similar patterns of ventricular enlargement ^51^ and hemisphere asymmetry ^54^ are reported. As a result, this model is suitable for investigating potential neuroprotective or reparative therapies that aim to prevent chronic structural changes after severe ischemic insults, such as neonatal strokes.

Consistent with ischemic insult causing persistent injury, we found that a marker of apoptotic cell death, CC3, was increased at 24 hours and remained elevated up to 14 days after injury. Elevated levels of cell death have been seen for at least 12 weeks after stroke in an adult rat model ^55^ and in adult models of traumatic brain injury ^56^. However, although CC3 is widely regarded as a key apoptotic executioner, emerging evidence highlights its non-apoptotic roles, including involvement in reactive astrogliosis and macrophage infiltration ^57^. Increases in CC3 immunoreactivity may relate to ongoing neuronal or microglial apoptosis ^58^ or as a specific inducer of a classically pro-inflammatory immune response in microglia in the region ^59^.

Effective therapies for severe focal brain injury in the neonate must modulate the neuroinflammatory response, given its central role in injury progression ^60–62^, including the morphological transition and increased expression of ionised calcium-binding adaptor molecule 1 (Iba1) by microglia and macrophages and GFAP-positive glial scar formation ^22,25,63–67^. In our model, GFAP peaked at 7 days after injury and declined by 14 days but remained significantly elevated compared to control levels. These findings are consistent with observations in models of severe neonatal focal brain injury, including HIE, neonatal stroke, or paediatric TBI in mice ^63^ ^68^, rats ^25,69^, non-human primates ^70^, piglets ^71^, and sheep^66^. Evidence suggests that the post-injury gliotic scar is less inhibitory to innate regeneration in the neonate. For instance, the post-stroke infant primate neocortex forms a smaller, more discrete chronic scar than in adults, correlating with greater neuronal sparing ^70^. Pro-regenerative neonatal astrocytes are also observed after injury to the neonatal spinal cord ^72^, setting the stage for further work to examine the phenotypic profile of these GFAP-reactive cells.

Ongoing microgliosis is another hallmark of perinatal brain damage and a potential target for the development of effective neurotherapeutics ^61,73^. In our model, macrophage/microglial density within the lesion core was dramatically elevated for at least 14 days, even as the lesion was reducing in volume. This could suggest a decoupling between tissue loss and inflammatory resolution, or that microglia are key in facilitating repair. Supporting the latter is that depletion of microglia before injury in a rat MCAO model of neonatal stroke increased pro-inflammatory markers and lesion volume ^74^, but the role of microglia is complex ^75^. This highlights that future studies should assess microglial phenotypic shifts in this model to determine whether they are predominantly deleterious or reparative, or whether function is context- and time-dependent^61,76,77^.

The present study found that higher doses of Rose Bengal increased the severity of brain damage, but this did not translate into a proportional increase in motor and sensory impairments, outcomes relevant to patients ^78^. Similar discrepancies between lesion volume and behavioural outcomes have been observed in young adult mice following photothrombotic stroke ^21,79^. In contrast, adult rats subjected to photothrombotic lesions displayed more pronounced neurological impairments and larger lesions ^80^. Similar variability has been observed in studies of the MCAO model, likely due to variability in the lesion territory ^80^, a limitation minimised in this model. However, we propose that the behavioural tests used may have limited sensitivity above a threshold of injury due to lesion cross-over into other regions, compensatory effects, or changes in mood behaviours that affect interactions with the behavioural paradigms ^81^. This may have masked subtle dose-dependent effects. Future studies should incorporate quantitative sensitive assays, such as automated gait analysis or a balance beam, to expand our understanding of the relationship between lesion territory, size, and functional outcomes.

We found no significant sex differences in lesion volume, behavioural performance, or inflammatory responses in this model. This contrasts with clinical and most preclinical studies that report greater injury severity, higher mortality rates, and more severe sensory-motor impairments in neonatal males than females after ischemic injury and in response to treatments ^38,82,83^, including in an adult model of photothrombotic infarct in the mouse ^84^. Our power calculations suggest that the effect sizes were large only for lesion volume at 24 hours; even then, to achieve statistical significance, 29 rats per group would have been required, and the group sizes for the other analyses were inappropriate to consider. This suggests that, under our experimental conditions, sex effects are biologically minimal. Further studies with longer follow-up periods and detailed analyses (e.g., transcriptomic or circuit-level approaches) are required to fully assess the impact of sex on outcomes in this model.

Histological and behavioural assessments were conducted up to 14 days post-injury, which poses a potential limitation, as this period is equivalent to 2-3 years in humans ^85,86^. A longer follow-up would provide a more comprehensive understanding of both recovery and chronic pathology. Second, while the behavioural tests employed detected major sensorimotor impairments, more challenging tasks, such as the rotarod, staircase test, or complex wheel, may better capture subtle or higher-order functional deficits. Third, we did not include detailed molecular profiling of glial subtypes or phagocytic microglial/macrophage activation states, which could offer deeper insights into the inflammatory and reparative mechanisms at play.

Despite the limitations, the model provides key practical advantages: a simple, fast and minimally invasive procedure with excellent long-term survival, high reproducibility, scalable lesion size, and measurable neurofunctional deficits. The consistency of lesion size and territory also enables efficient parallel testing of candidate therapeutics while maintaining animal welfare, making it a valuable tool for studies of severe neonatal focal brain injury.

## CRediT (Contributor Roles Taxonomy) statement

**Conceptualization**: V.M.; E.R-M.; J.L.Y.C.; F.Y.W.; D.W.W.; M.T.; B.F.; **Methodology**: V.M.; E.R-M.; G.K.B.; R.K.; I.K.S.; A.M.; A.A.M.S.; M.B; P.G.; J.L.Y.C.; F.Y.W.; D.W.W.; M.T.; B.F.; **Formal analysis**: V.M.; A.A.M.S.; M.B.; D.W.W.; M.T.; B.F.; **Data Curation**: V.M.; E.R.M.; G.K.B.; R.K.; I.K.S.; A.M.; A.A.M.S.; M.B.; D.W.W.; M.T.; B.F.; **Writing - Original Draft; Writing:** V.M.; B.F.; **Writing - Review & Editing**: V.M.; E.R.M.; G.K.B.; R.K.; I.K.S.; A.M.; A.A.M.S.; M.B.; D.R.N.; G.F.K.; R.J.W.; P.G.; J.L.Y.C.; F.Y.W.; D.W.W.; M.T.; B.F.; **Supervision:** E.R.M.; G.K.B.; I.K.S.; M.B.; D.R.N.; G.F.K.; R.J.W.; D.W.W.; M.T.; B.F.; **Project administration**: V.M.; M.T.; B.F.; **Funding acquisition** D.R.N.; G.F.K.; R.J.W.; P.G.; J.L.Y.C.; F.Y.W.; D.W.W.; M.T.; B.F.

## Supporting information

Summ methods, figrues and tables

## Acknowledgments

We acknowledge funding support from the Cerebral Palsy Alliance (B.F., D.W.W., and M.T.), the Kinghorn Foundation (B.F., D.W.W., G.F.K., R.W., and M.T.), the National Health and Medical Research Council of Australia (APP1187607 to B.F., APP1161466 to M.T. and D.W.W., Investigator Fellowships 2035090 to G.F.K. and 2016390 to J.L.Y.C.), The Lott by Golden Casket (G.F.K.), and the Bruce Hyams Rising Star Fellowship (K.F.). This work was also supported by the Australian Research Council via Future Fellowships FT230100220 (D.N.) and FT180100082 (M.T), and the ARC Centre for Innovations in Peptide and Protein Research (CE200100012). and to P.G support was provided from Inserm, Inserm IRP, Université Paris Cité, Horizon 2020 Framework Program of the European Union (grant agreement no. 874721/PREMSTEM), ANR, ERANET-NEURON (VasOx), DIM C-Brains (Ile de France region), QIM Veave (Ile de France region), Fondation de France, Fondation pour la Recherche sur le Cerveau, Fondation Grace de Monaco, Fondation des Gueules Cassées, ANR-23-IAHU-0010 (France 2030), and an additional grant from “Investissement d’Avenir-ANR-11-INBS-0011-“ NeurATRIS.

