## Supplementary material for "A Minimally Invasive, Scalable and Reproducible Neonatal Rat Model of Severe Focal Brain Injury": Summ methods, figrues and tables

**Supplementary figures**

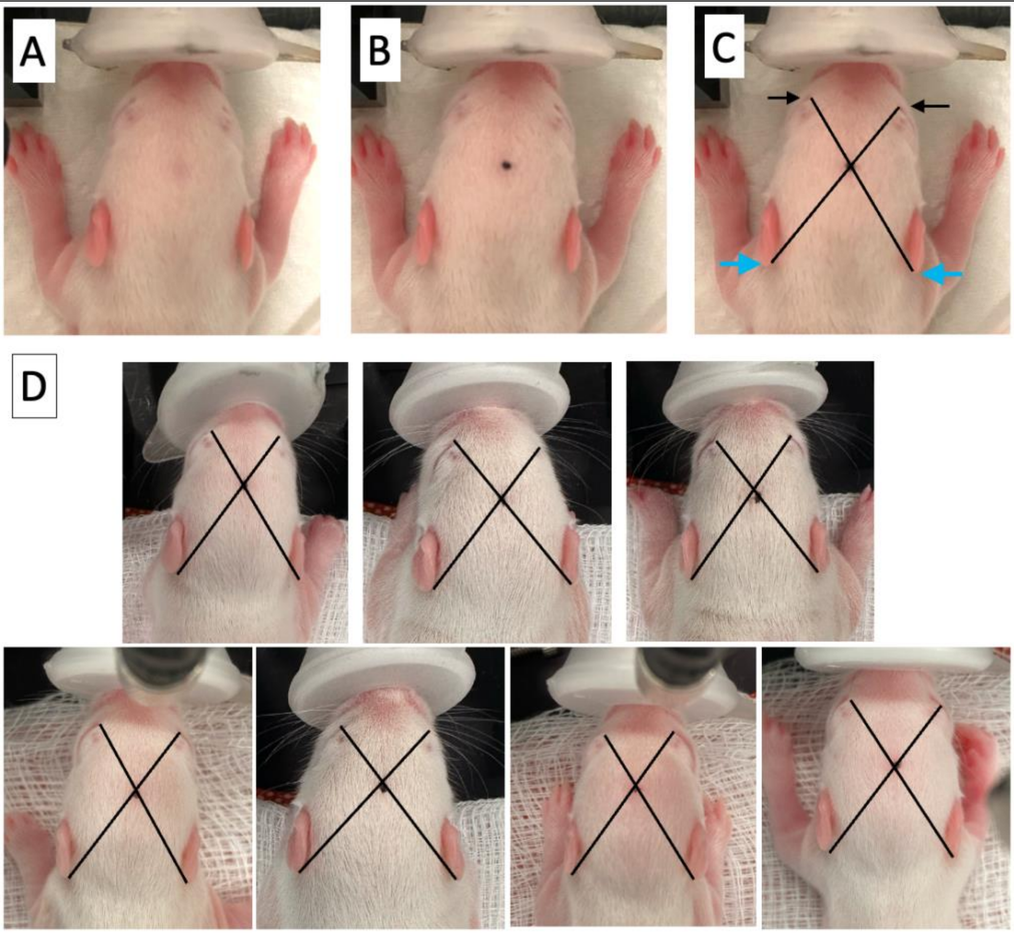


**Supplementary Figure 1.** The view onto the top of the rat pups head, showing the (**A**) reliably present darker patch of hair that can be (**B**) used for marking our bregma which (**C**) lays on a diagonal line between the medial canthus (inner eye crease, black arrows) and the posterior border of the ear lobe (blue arrows) when viewed from above. (**D**) Examples of the location of the bregma relative to the anatomical regions from across experiments.

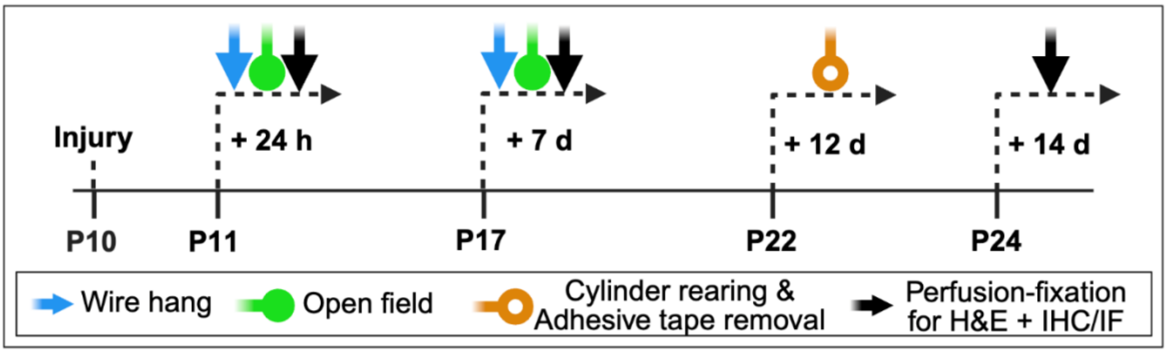


**Supplementary Figure 2.** Experimental design outline. *Abbreviations*: d, day; h, hour; H&E, hematoxylin and eosin; IF, immunofluorescence; IHC, immunohistochemistry.

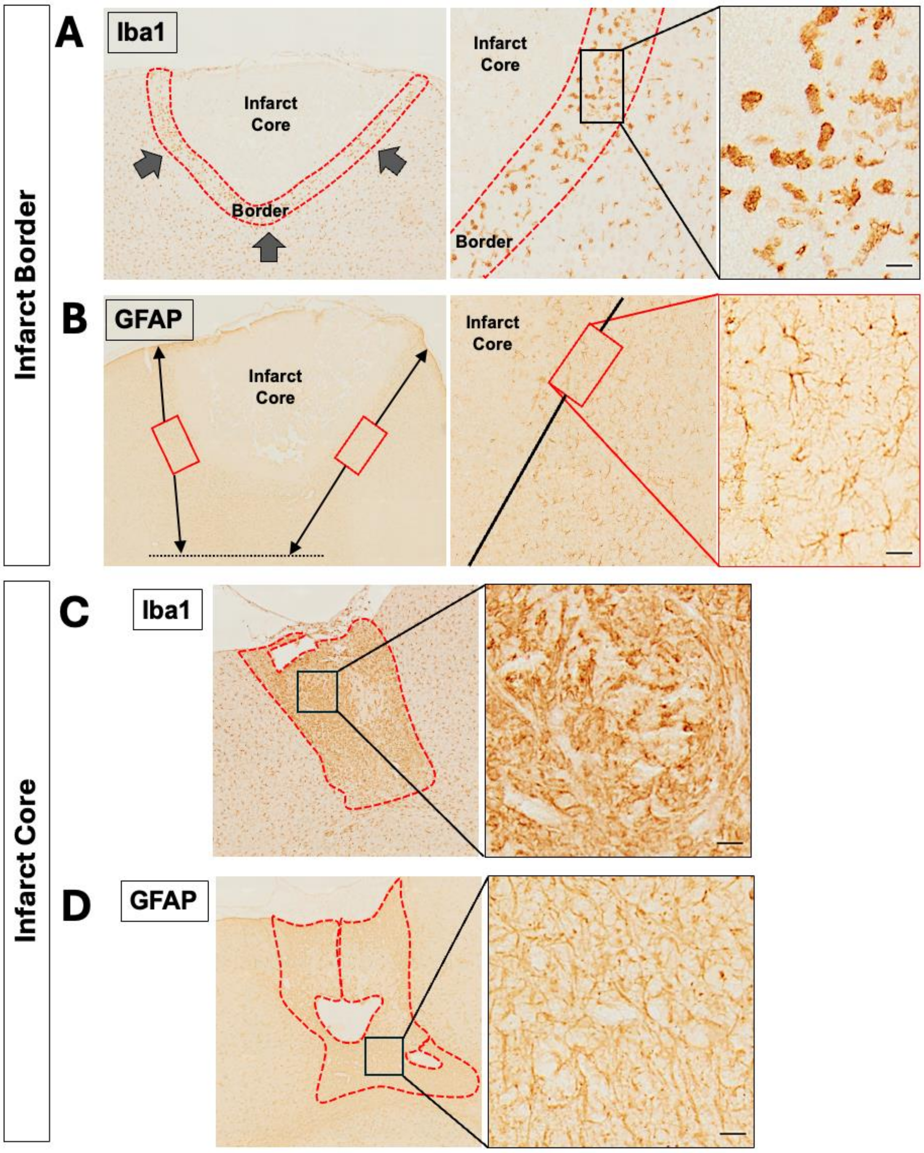


**Supplementary Figure 3. Explanation of the locations of the zones of interest for Iba1 and GFAP staining analyses.** In (**A**), the polygon tool was used to trace around the band of ameboid Iba1-positive cells present at the border of the infarct at 24 hours, referred to as the infarct border. Also shown is an inset of the infarct border zone with arrows to mark its width (upper left) and then higher magnification insets to show inside the border zone and the morphology of Iba1-positive cells (upper right). In (**B**), a representative image of GFAP staining at 24 hours, showing the placement of the two fixed-size regions of interest (ROIs) in the lesion by drawing straight lines from the upper cortical injury border to the lower injury border. The ROIs were placed halfway between the upper and lower infarct boundaries. A higher magnification inset shows inside the GFAP infarct border zone and the morphology of GFAP-positive cells (lower right). In (**C**) the infarct core zone is present at all timepoints analysed in Iba1 (upper) and in (**D**) GFAP (lower) stained sections. The infarct core zone was delineated using the polygon tool on a low-magnification overview of the lesion, characterised by an area densely populated with microglia/macrophages and astrocytes (left). Higher magnification insets (right panel) illustrate the reactive glial cells within the infarct core. Scale bars = 20 μm.

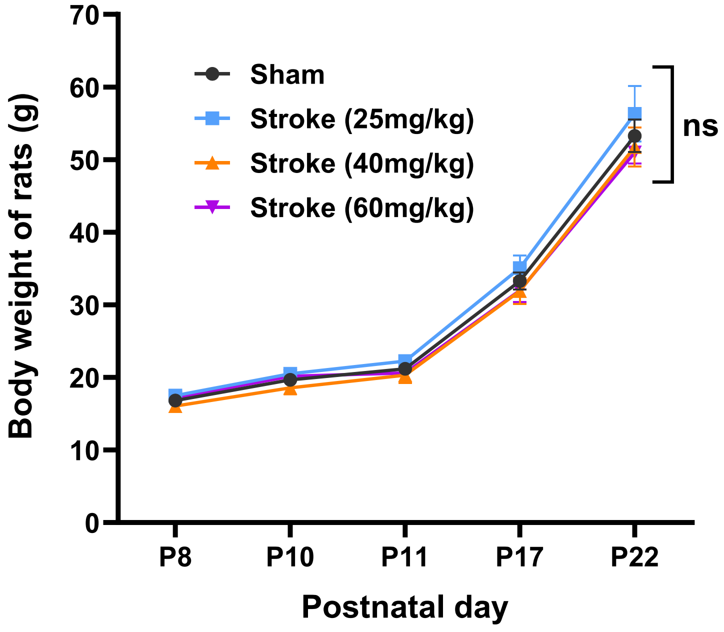


**Supplementary Figure 4.** Weight gain for sham and the three Rose Bengal groups (mean ± SEM). Statistical analysis was performed with a two-way ANOVA followed by Sidak's *post-hoc* multiple comparison test. ns = not significant.

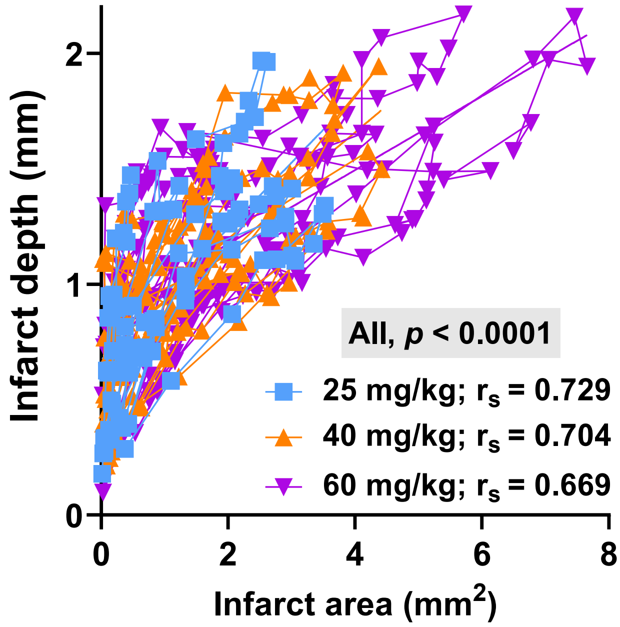


**Supplementary Figure 5.** Linear correlations between infarct area and infarct depth by dose, with Spearman's correlation coefficient ($r_s$) and p-values shown.

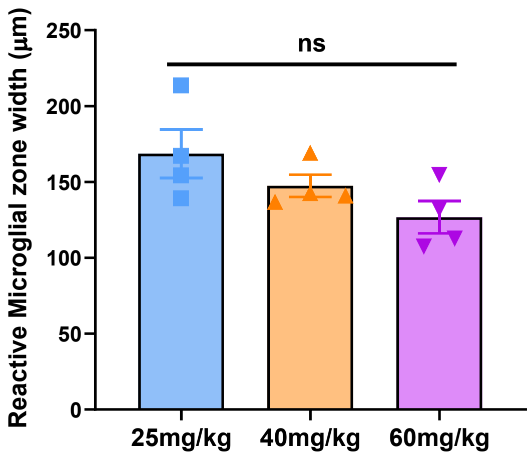


**Supplementary Figure 6.** Ameboid Iba1-IR zone width measurements at 24 hours post-stroke. n = 4 for each dose. Data are mean ± SEM, ns = not significant from one-way ANOVA, Sidak's *post-hoc* analysis.

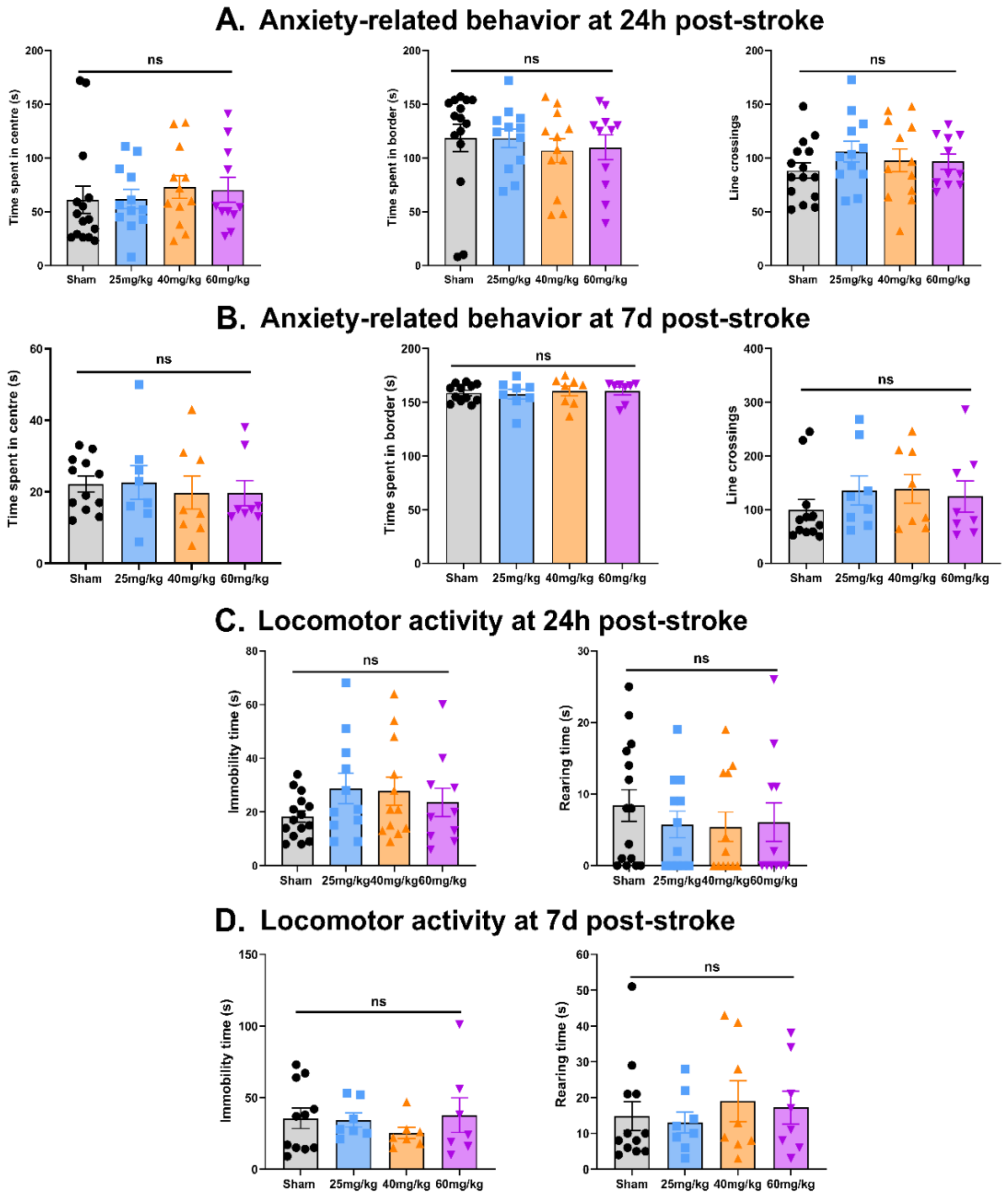


**Supplementary Figure 7.** Assessment of the behavioural changes in the open field test following neonatal stroke at 24 hours and 7 days post-stroke. Sham (n = 15), 25 mg/kg (n = 12), 40 mg/kg (n = 12), 60 mg/kg (n = 11) at 24h post-stroke and Sham (n = 12), 25 mg/kg (n = 8), 40 mg/kg (n = 8), 60 mg/kg (n = 8) at 7d post-stroke. Data are mean ± SEM, ns = not significant from one-way ANOVA, Sidak's *post-hoc* analysis.

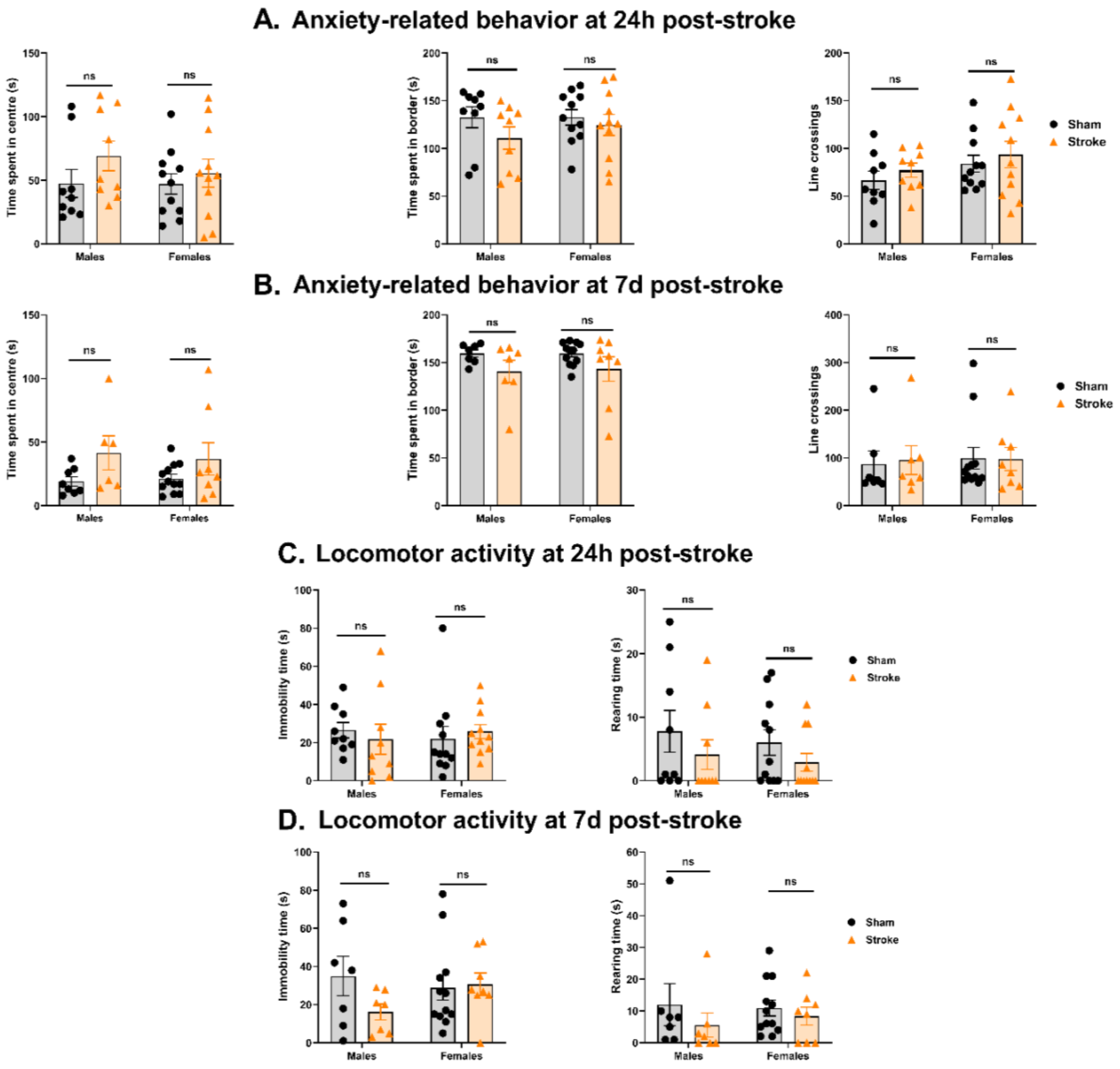


**Supplementary Figure 8.** Assessment of the behavioural changes in the open field test following neonatal stroke at 24 hours and 7 days post-stroke in males and females. At 24 hours post-stroke, sham males (n = 9), sham females (n = 11), stroke males (n = 9), stroke females (n = 11), at 7 days, sham males (n=8), sham females (n = 12), stroke males (n = 6), stroke females (n = 8) post-stroke. Data are mean ± SEM, ns = not significant from Two Way ANOVA, Sidak's *post-hoc* analysis.

### Materials and methods

#### Animals and surgical procedures

All experiments were approved by the RMIT University Animal Ethics Committee (AEC 1918), and work is reported in alignment with the ARRIVE guidelines. Rats were obtained from Australian Bioresources (Perth, Western Australia) at 12–16 days of pregnancy and allowed to deliver undisturbed. Body weights were recorded for all animals on postnatal (P) day 8, and each animal was assigned a random identifier. Then, using a restricted randomisation approach, they were group-allocated within litters in a block design. P10 was chosen for injury induction because it is considered approximately equivalent to a term-born human infant in terms of brain development [32]. Predetermined exclusion criteria included situations in which animals needed to be euthanised prior to planned experimental endpoints for humane reasons, or technical issues with tissue processing or staining. Only one animal died before the assigned collection point (sham group), and no other animal was excluded at any point across the analyses. For neuropathological assessments, one animal per litter per time point was included. For behaviours, each pup in the litter was tested and included as an independent replicate. For sex analysis, additional animals were added to the 25 mg/kg group.

P10 male and female Wistar rats were anaesthetised initially with 5% isoflurane (Isoflo, Abbott, NSW, Australia) in 0.4 L/minute oxygen by inhalation from a face mask for 2 minutes, then anaesthesia was maintained with 2–3% isoflurane in 0.4 L/minute oxygen. Body temperature was maintained using a thermistor-controlled heating pad (Thorlabs, USA). Rose Bengal (Merck, Australia, cat#330000), dissolved in sterile saline solution and protected from light was injected intraperitoneally at 25, 40 or 60 mg/kg, and after 5 minutes, a 1 mm diameter laser emitting diode (LED) light source (Thorlabs, US, cat# M565F3, 565 nm) was placed on the skin 2 mm to the right of Bregma and illuminated on full power for 10 minutes to induce a photothrombotic cortical injury. Doses of Rose Bengal were selected to span a range of possible injury severities, based on a review of the literature[25].

When viewed from above, Bregma appears as a dark patch at the intersection of an imaginary line drawn between the medial canthus (inner ear crease) and the posterior border of the ear lobe when viewed from above (**Supplementary Figure 1**). At the end of the surgery, which took on average 19 minutes, buprenorphine (0.05 mg/kg; stock 300 μg/mL in sterile saline, Cenvet, Australia) and meloxicam (5 mg/kg; stock 5 mg/mL in sterile saline, Cenvet, Australia) were injected subcutaneously, and after a further 60 minutes of recovery on a pre-warmed heating pad, pups were returned to the dam. Sham controls were administered Rose Bengal without laser illumination to account for the confounding effects of the dye, which can affect the vasculature [33]. Light exposure has no reported discernible effects and was not considered a confounding factor.

**Behavioural testing**

Behavioural testing was undertaken between 0800 and 1200 hours. Rats were identified only via the number allocation from P8. Rats were individually transferred from their home cage to a holding cage, then to the testing area. All behavioural equipment was thoroughly cleaned with 70% ethanol and allowed to dry to eliminate residual odours between test sessions. The testing scheme is summarised in **Supplementary Figure 2**. P11 (24 hours post-injury) was selected to capture the acute phase of neuroinflammation and glial activation, corresponding to a stage in rodent brain development broadly equivalent to a human term infant. P17 (7 days post-injury) reflects the subacute phase, during which lesion consolidation and active remodelling occur, aligning with the end of the neonatal period in humans. P24 (14 days post-injury) represents the early recovery phase, coinciding with a surge in myelination and cortical circuit maturation in rats, developmentally analogous to late infancy to early toddlerhood (approximately 2–3 years in humans) [3234,35].

*Wire hang test.* Wire hang tests were conducted at P11 and P17. The rats were held by the scruff of the neck skin and supported until they grasped the wire with both forepaws. They were allowed to hang for a maximum of 5 minutes. Three separate hang times were recorded for each rat, with a 30-second rest period between each trial. Trials were recorded, and the total time spent hanging on the wire was logged from the video footage. The distance from the wire to the surface below was approximately two body lengths at P11 (i.e., 24 hours after inducing the injury), and approximately three body lengths at P17 (7 days after the injury).

*Cylinder rearing test.* At P22 (i.e., 12 days post-injury), rats were placed in a transparent plexiglass cylinder (185 mm height x 130 mm outer diameter). Video footage was recorded for 5 minutes using a camera positioned beneath the cylinder, capturing an upward view of the rat. The number of times the rat reared onto its hindlimbs and contacted the cylinder with one or both forepaws was recorded. Wall touches were categorised and counted separately for each forepaw (right and left) as well as for simultaneous bilateral contacts (double contact). Data were expressed as a percentage of the impaired forelimb touches relative to total forelimb contacts, calculated as: $= \left(\frac{(contralateral\ touches)}{(ipsilateral\ touches + contralateral\ touches)}\right) \times\ 100$ .

*Adhesive tape removal test.* At P22, two identical pieces of adhesive tape (Tough-Spots, 0.95 cm diameter, halved, Merck, USA, Z743588) were affixed to the hairless area of the palm of each forepaw. Rats were immediately placed in a transparent plexiglass cylinder (185 mm height x 130 mm outer diameter) with video recording from underneath. Each test session consisted of three trials, with a 1-minute rest period between each. Each trial had a maximum duration of 3 minutes; if the tape was not removed within this period, the trial was recorded for up to three minutes. Initial contact time was defined as the first instance of paw shaking or bringing the paw to the mouth. Sensorimotor performance was assessed by measuring the time taken to remove the adhesive from each paw.

*Open field test.* At P11 and P17, rats were placed in the centre of a square arena (30 x 30 cm) and allowed to explore freely for 3 minutes, recorded via an overhead camera. Parameters analysed included horizontal activity (number of line crossings), time spent in the centre and periphery zones, immobility duration, and rearing behaviour.

**Brain tissue collection and processing**

Brains were collected from injured and sham rats at P11, P17, and P24 (**Figure 1**). Rats were anesthetised with an intraperitoneal injection of Lethabarb (325 mg/mL solution, >200 mg/kg, Virbac, Australia, plus 10 mg/mL Lignocaine, Cenvet, Australia) and transcardially perfused with 0.1 M phosphate buffered saline (PBS, pH 7.4) for 1 minute followed by 4% paraformaldehyde (PFA, Sharlau, Barcelona, Spain), for 4 minutes at a flow rate of approximately 2 mL/minute. Brains were post-fixed for 12-18 hours at 4°C in a 1:10 volume of PFA, then transferred to 70% ethanol. Tissues were processed (Leica Microsystems, Australia, model ASP300S) and embedded in paraffin wax in a coronal orientation. Processing conditions were 1 hour each in the following: 70% ethanol, 90% ethanol, 100% ethanol, 50/50 ethanol/xylene, xylene, fresh xylene, paraffin wax, fresh paraffin wax, and fresh paraffin wax. Coronal sections were cut on a rotary microtome (Thermoline Scientific, Australia, model HM325), starting at approximately Bregma +3.24, before the lesion appeared, and continuing through the brain until no lesion was present. Sections were cut at 8 µm, transferred to float in a 37°C water bath, and every twentieth section (three sections per slide, spaced 160 µm apart) was mounted onto Superfrost Plus slides (Menzel-Gläser, Germany) and air-dried before storage.

### Infarct, cortical atrophy and ventricle asymmetry analysis

*Haematoxylin and Eosin (H&E) staining*. Every twentieth section throughout the lesion was stained. The slides were deparaffinized, rehydrated, stained in 0.3% Mayer's Haematoxylin (Amber Scientific, Australia) for 4 minutes, washed in running tap water until the water was colourless, and then submerged in Scott's tap water (20 g of magnesium sulphate and 2 g of sodium bicarbonate, both from Sigma Aldrich, in 1 litre of deionized water) for 1 minute. Sections were then rinsed in running tap water again for 3 minutes, counterstained with 1% alcohol eosin (Amber Scientific, Australia) for 3 minutes, dehydrated and cover-slipped with dibutylphalate polystyrene xylene (DPX; Merck, Australia).

*Infarct volume, location and depth analysis.* Images were collected at x10 magnification using the Olympus VS120 Virtual Slide Microscope (Olympus Life Science, VIC, Australia) All analyses were performed by experimenters blinded to group assignments. The area and depth of the infarct were measured using ImageJ (ImageJ 1.53c Wayne Rasband, National Institutes of Health, USA) as previously[36,37]. The total infarct volumes were estimated using Cavalieri's principle by the following formula: $V = SA \times P \times T$, where V is the total volume, SA is the sum of areas measured, P is the inverse of the sampling fraction (20), and T is the section thickness (8 µm).

According to the Paxinos and Watson rat brain atlas [38], the area of the lesion from each section was plotted against its location relative to Bregma to assess the reproducibility of lesion location. The infarct area within the primary motor cortex was calculated using this formula: $= \left(\frac{Infarct\ area\ located\ in\ the\ primary\ motor\ cortex}{Total\ infarct\ area}\right) \times 100$. Infarct depth was measured from the mid-point of the lateral-to-medial edge of the lesion on the outermost cortical surface to the deepest visible lesion using the line tool in ImageJ.

The reproducibility of the lesion between litters of rats was assessed by calculating the coefficient of variation (CV) [39], as $\left(\frac{standard\ deviation}{mean}\right) \times 100$ for the lesion volume ($CV_{lesion}$), and the lesion area as a percentage of the hemisphere ($CV_{percent}$), expressed as a percentage.

*Analysis of cortical atrophy and ventricle asymmetry.* Brain atrophy in the affected right hemisphere was quantified by measuring the area of every 20th H&E-stained section and comparing it to the contralateral hemisphere. Cortical atrophy was calculated for each section using the formula: Cortical atrophy (%) = $\left[1 - \left(\frac{ipsilateral\ area}{controlateral\ area}\right)\right] \times 100$, representing the percentage of ipsilateral hemisphere atrophy, as previously [37,40]. Values were averaged across all analysed sections. Lateral ventricle size was also measured, and the lateral ventricle ratio (LVR) was calculated as the ipsilateral to contralateral lateral ventricle area ratio using ImageJ.

**Immunohistochemistry (IHC) and immunofluorescence (IF)**

Coronal sections spaced 160 µm through the lesion were dewaxed, rehydrated, and subjected to antigen retrieval as previously described [41,42]. All sections to be compared were processed within the same run to prevent inter-batch variation. Antigen retrieval was performed in citrate buffer (pH 6.0) heated to boiling in a microwave (Whirlpool M542), followed by two 7-minute heating cycles at 80% power with a 5-minute interval, and cooling in buffer for 20 minutes.

For IF, sections were washed in PBS (3 × 5 minutes), incubated in 0.1% Sudan Black B (Abcam, MA, USA; 0.1% in 70% ethanol) for 20 minutes to reduce autofluorescence, washed again, and blocked in 5% normal goat serum (NGS; ThermoFisher, Australia, cat# 10000C) with 0.2% Triton X-100 (Sigma-Aldrich, Australia, cat# 10789704001) in PBS for 1 hour at room temperature. For IHC, endogenous peroxidase activity was quenched with 3% $H_2O_2$ (Chem-Supply, Australia, cat# HA154-2.5L-P), then sections were blocked with 5% NGS and 0.2% Triton X-100 in PBS for 1 hour.

Sections were incubated overnight at 4 °C in a humidified chamber with the following primary antibodies diluted in 1% NGS/0.1% Triton X-100: anti-GFAP (rabbit polyclonal, 1:1000; Dako, cat# Z033401-2), anti-Iba1 (rabbit polyclonal, 1:1000; Novachem, cat# 019-9741), and anti-cleaved caspase-3 (rabbit polyclonal, 1:500; Cell Signalling, cat# 966L1).

The following day, IF sections were washed and incubated for 1 hour at room temperature with donkey anti-rabbit Alexa Fluor 488 secondary antibody (Invitrogen, cat# A32790), followed by DAPI nuclear counterstaining (ThermoFisher, USA; cat# D2149; 1:1000, 15 minutes). IHC sections were incubated for 1 hour with goat anti-rabbit biotinylated secondary antibody (Vector Laboratories, cat# VEBA1000), followed by Vectastain ABC Elite (Vector Laboratories, cat#

PK6100; 1:1:200) and visualisation with DAB (50 mg in 10 mL $dH_2O$ plus 3 µL 30% $H_2O_2$). All slides were washed thoroughly between steps (3 × 5 minutes).

IF sections were mounted in Vectashield aqueous medium (Vector Laboratories, USA) and protected from light. IHC slides were dehydrated and coverslipped with DPX. Slides dried for 24–48 hours before imaging.

**Image analysis**

*Image acquisition and processing.* Images were acquired using an Olympus VS120 slide scanner at x10 magnification for H&E and at x20 for IHC and IF. The exported data were analysed using ImageJ (ImageJ 1.53c Wayne Rasband, National Institutes of Health, USA), Excel (Microsoft 365, USA), and Prism (version 8, GraphPad Software Inc., USA).

*Quantification of immunohistochemically stained areas.* All analyses were performed blinded to group allocation. Pilot experiments at 6, 24, and 48 h and at 7 and 14 d post-injury were used to determine optimal time points and regions of interest (ROIs; **Supplementary Figure 3**). For each marker, three coronal sections were analysed, at the lesion's maximal extent and one-third either side.

At 24 h post-injury (P11), Iba1-positive microglia/macrophages were quantified in the infarct border (polygonal ROI), infarct core (polygonal ROI), and a fixed-size ROI in the corresponding sham cortex. At 7 and 14 days (P17, P24), Iba1 was assessed in the infarct core and sham cortex. GFAP-positive astrocytes and CC3-positive apoptotic cells were quantified at P11 in two fixed-size ROIs (200 µm × 400 µm) positioned laterally along the astrocyte reactivity zone, within the infarct core, and in the sham cortex; at later time points, these markers were analysed in the core and sham regions only.

The infarct border was defined as tissue surrounding the core between 24 and 48 h post-injury, whereas the core itself showed grossly altered morphology, acellular at early stages and densely glial at later stages. Astrocytic scar formation and microglial/macrophage accumulation were delineated using the polygon tool in ImageJ, as illustrated in **Supplementary Figure 3**. The percentage area of Iba1- and GFAP-positive staining was determined by applying a consistent pixel-intensity threshold to 8-bit converted images. Background intensity was calculated from representative samples across groups, averaged, and then subtracted uniformly. The proportion of positive pixels was expressed as the number of thresholded pixels within the ROI relative to the total ROI area. CC3-positive cells were counted within the same ROIs used for GFAP analysis.

### Statistical analyses

Data are presented as mean ± standard error of the mean (SEM). Statistical analyses were performed using Prism (version 8; GraphPad Software Inc., USA). A p-value $< 0.05$ was considered statistically significant. Parametric tests were used unless otherwise stated, based on their robustness to modest deviations from normality and the limited power of non-parametric alternatives in small *n* datasets. Infarct volume, cortical atrophy, and ventricular enlargement were analysed using two-way ANOVA followed by a Sidak's post-hoc multiple comparisons test. The relationship between infarct area and depth was assessed using Spearman's rank correlation coefficient (ρ), with corresponding $r_s$ and p values reported. Differences in total wire hang time, cylinder rearing time, and adhesive tape removal time between the sham and injury groups were analysed using one-way or two-way ANOVA with Sidak's post-hoc multiple comparisons when a significant interaction or main effect was observed, as indicated in the figure legends. The number of cylinder wall touches by forepaws was analysed using two-way ANOVA with Sidak's post-hoc test, where there was a significant interaction or main effect. CC3, Iba1 and GFAP staining in the infarct border were assessed using one-way ANOVA, while staining in the infarct core was analysed using two-way ANOVA with Sidak's multiple comparisons, as indicated in the figure legends. Correlations between behavioural outcomes and lesion volumes were assessed using Pearson's correlation coefficient. Sex-specific analyses were performed using two-way ANOVA with Sidak's post-hoc test. ANOVA main effects are reported in the **Supplementary Tables**, and the results of post-hoc tests are shown in the figures. Power analyses were performed using G*Power (version 3.1) [43].

### Supplementary tables

**Supplementary Table 1.** Comparison of weight gain between sham and focal injury rat pups at postnatal days

| | **P10** | **P11** | **P17** | **P22** |
|---|---|---|---|---|
| **Sham pup no., (*n*, *litters*)** | 18 (*4*) | 18 (*4*) | 13 (*4*) | 9 (*4*) |
| **Average sham body weight (g)** | 19.6 ± 0.6 | 21.2 ± 0.6 | 33.3 ± 1.2 | 53.3 ± 2.3 |
| **25 mg/kg injury group average body weight in gram (*pup n, litter n*)** | 20.5 ± 0.7 (*n=12, 4*) | 22.2 ± 0.7 (*n=12, 4*) | 35.1 ± 1.7 (*n=8, 4*) | 56.3 ± 3.8 (*n=4, 4*) |
| **Post-hoc p-value 25 mg/kg RB vs sham** | 0.878 | 0.790 | 0.578 | 0.366 |
| **40 mg/kg injury group average body weight in gram (*pup n, litter n*)** | 18.6 ± 0.9 (*n=12, 4*) | 20.3 ± 1.1 (*n=12, 4*) | 31.9 ± 1.8 (*n=8, 4*) | 51.7 ± 2.7 (*n=4, 4*) |
| **Post-hoc p-value 40 mg/kg RB vs sham** | 0.782 | 0.872 | 0.749 | 0.841 |
| **60 mg/kg injury group average body weight in gram (*pup n, litter n*)** | 20.2 ± 0.7 (*n=12, 4*) | 20.6 ± 0.8 (*n=12, 4*) | 32.0 ± 1.6 (*n=8, 4*) | 51.0 ± 1.5 (*n=4, 4*) |
| **Post-hoc p-value 60 mg/kg RB vs sham** | 0.973 | 0.965 | 0.789 | 0.602 |

Body weights (mean ± SEM) and p-values are from a two-way ANOVA, followed by Sidak's-adjusted multiple comparison test. **Abbreviations:** g, gram; n, number; P, postnatal day; RB, Rose Bengal; vs, versus.

**Supplementary Table 2. Reproducibility data for lesion volume at 24 hours after injury**

| | **25 mg/kg** | | **40 mg/kg** | | **60 mg/kg** | |
|---|---|---|---|---|---|---|
| | Lesion Vol. | Lesion Area (% Hemi.) | Lesion Vol. | Lesion Area (% Hemi.) | Lesion Vol. | Lesion Area (% Hemi.) |
| **Mean** | 3.74 | 8.15 | 5.33 | 8.36 | 11.15 | 11.76 |
| **SD** | 1.16 | 2.20 | 1.07 | 1.81 | 1.27 | 1.85 |
| **CV (%)** | 31% | 27% | 20% | 22% | 11% | 16% |

**Data are presented as mean ($mm^3$) ± SD (*n* = 4/dose). Abbreviations: CV, coefficient of variation; Hemi., hemisphere; SD, standard deviation; Vol., volume.**

**Supplementary Table 3.** Results of the statistical analysis for infarct volume comparisons

| Figure | Test name | Main effects | | | Sidak's multiple comparison | | | |
|---|---|---|---|---|---|---|---|---|
| | | Interaction F (DFn, DFd), p value | Stroke doses, F (DFn, DFd), p value | Time post-stroke, F (DFn, DFd), p value | Time post-stroke | Stroke doses comparisons | 95% CI | p value |
| **Figure 1B,** Infarct volume | Two-way ANOVA | F (4, 27) = 10.43, ***p < 0.0001*** | F (2, 27) = 40.78, ***p < 0.0001*** | F (2, 27) = 106.1, ***p < 0.0001*** | 24 h | 25 mg/kg vs 40 mg/kg | -3.492 to 0.3142 | 0.1233 |
| | | | | | | 25 mg/kg vs 60 mg/kg | -9.321 to -5.515 | ***< 0.0001*** |
| | | | | | | 40 mg/kg vs 60 mg/kg | -7.732 to -3.927 | ***< 0.0001*** |
| | | | | | 7 d | 25 mg/kg vs 40 mg/kg | -2.898 to 0.9080 | 0.4772 |
| | | | | | | 25 mg/kg vs 60 mg/kg | -3.881 to -0.07525 | ***0.0397*** |
| | | | | | | 40 mg/kg vs. 60 mg/kg | -2.886 to 0.9197 | 0.4872 |
| | | | | | 14 d | 25 mg/kg vs 40 mg/kg | -2.643 to 1.163 | 0.7011 |
| | | | | | | 25 mg/kg vs 60 mg/kg | -3.880 to -0.07413 | ***0.0399*** |
| | | | | | | 40 mg/kg vs 60 mg/kg | -3.140 to 0.6654 | 0.2938 |

Infarct volume data were analysed using two-way ANOVA to compare three Rose Bengal doses at multiple timepoints, with significance set at $p < 0.05$. Sidak's post-hoc test was used for multiple comparisons. **Bold and italicised values indicate statistically significant differences.** Sample sizes: 25 mg/kg = 4 rats, 40 mg/kg = 4 rats, 60 mg/kg = 4 rats. **Abbreviations:** ANOVA, analysis of variance; CI, confidence interval; DFd, degrees of freedom (denominator); DFn, degrees of freedom (numerator); *p*, probability value.

**Supplementary Table 4.** Results of the statistical analysis for cortical atrophy and ventricle enlargement

| Figure | Test name | Main effects | | | Sidak's multiple comparison | | | |
|---|---|---|---|---|---|---|---|---|
| | | Interaction F (DFn, DFd), p value | Stroke doses, F (DFn, DFd), p value | Time post-stroke, F (DFn, DFd), p value | Time post-stroke | Stroke doses comparisons | 95% CI | p value |
| **Figure 1D,** Cortical atrophy | Two-way ANOVA | F (6, 36) = 1.811, p = 0.1245 | F (3, 36) = 26.74, ***p < 0.0001*** | F (2, 36) = 9.881, ***p = 0.0004*** | 24 h | Sham vs 25 mg/kg | -9.479 to 0.8295 | 0.1420 |
| | | | | | | Sham vs 40 mg/kg | -9.677 to 0.6322 | 0.1122 |
| | | | | | | Sham vs 60 mg/kg | -11.57 to -1.263 | ***0.0083*** |
| | | | | | | 25 mg/kg vs 40 mg/kg | -5.352 to 4.957 | >0.9999 |
| | | | | | | 25mg/kg vs 60 mg/kg | -7.247 to 3.062 | 0.8435 |
| | | | | | | 40 mg/kg vs 60 mg/kg | -7.050 to 3.259 | 0.8947 |
| | | | | | 7 d | Sham vs 25 mg/kg | -14.96 to -4.656 | ***<0.0001*** |
| | | | | | | Sham vs 40 mg/kg | -12.81 to -2.504 | ***0.0012*** |
| | | | | | | Sham vs 60 mg/kg | -18.59 to -8.276 | ***<0.0001*** |
| | | | | | | 25 mg/kg vs 40 mg/kg | -3.002 to 7.306 | 0.8260 |
| | | | | | | 25 mg/kg vs 60 mg/kg | -8.775 to 1.534 | 0.3030 |
| | | | | | | 40 mg/kg vs 60 mg/kg | -10.93 to -0.6178 | ***0.0213*** |
| | | | | | 14 d | Sham vs 25 mg/kg | -9.663 to 0.6457 | 0.1141 |
| | | | | | | Sham vs 40 mg/kg | -11.91 to -1.599 | ***0.0050*** |
| | | | | | | Sham vs 60 mg/kg | -13.20 to - | ***0.0007*** |

| | | | | | | | | |
|---|---|---|---|---|---|---|---|---|
| | | | | | | | 2.892 | |
| | | | | | | 25 mg/kg vs 40 mg/kg | -7.399 to 2.910 | 0.7971 |
| | | | | | | 25 mg/kg vs 60 mg/kg | -8.692 to 1.617 | 0.3278 |
| | | | | | | 40 mg/kg vs 60 mg/kg | -6.448 to 3.861 | 0.9823 |
| **Figure 1E,** Ventricle enlargement | Two-way ANOVA | F (6, 36) = 1.087, p = 0.3884 | F (3, 36) = 12.22, ***p < 0.0001*** | F (2, 36) = 0.5672, p = 0.5721 | 24 h | Sham vs. 25 mg/kg | -22.72 to -2.906 | ***0.0057*** |
| | | | | | | Sham vs 40 mg/kg | -18.84 to 0.9762 | 0.0963 |
| | | | | | | Sham vs 60 mg/kg | -22.53 to -2.711 | ***0.0066*** |
| | | | | | | 25 mg/kg vs 40 mg/kg | -6.026 to 13.79 | 0.8637 |
| | | | | | | 25 mg/kg vs 60 mg/kg | -9.714 to 10.10 | >0.9999 |
| | | | | | | 40 mg/kg vs 60 mg/kg | -13.60 to 6.221 | 0.8894 |
| | | | | | 7 d | Sham vs 25 mg/kg | -15.00 to 4.814 | 0.6511 |
| | | | | | | Sham vs 40 mg/kg | -13.57 to 6.246 | 0.8925 |
| | | | | | | Sham vs 60 mg/kg | -23.47 to -3.651 | ***0.0031*** |
| | | | | | | 25 mg/kg vs 40 mg/kg | -8.476 to 11.34 | 0.9991 |
| | | | | | | 25 mg/kg vs 60 mg/kg | -18.37 to 1.444 | 0.1294 |
| | | | | | | 40 mg/kg vs 60 mg/kg | -19.81 to 0.011 | 0.0504 |
| | | | | | 14 d | Sham vs 25 mg/kg | -19.15 to 0.6687 | 0.0787 |
| | | | | | | Sham vs 40 mg/kg | -19.44 to 0.3812 | 0.0640 |

| | | | | | | | | |
|---|---|---|---|---|---|---|---|---|
| | | | | | | Sham vs 60 mg/kg | -19.46 to 0.3587 | 0.0987 |
| | | | | | | 25 mg/kg vs 40 mg/kg | -10.20 to 9.621 | >0.9999 |
| | | | | | | 25 mg/kg vs 60 mg/kg | -10.22 to 9.599 | >0.9999 |
| | | | | | | 40 mg/kg vs 60 mg/kg | -9.931 to 9.886 | >0.9999 |

Two-way ANOVA was used to analyse cortical atrophy and ventricle enlargement across three Rose Bengal doses and sham controls, with significance set at $p < 0.05$. Sidak's post hoc test was used for multiple comparisons. Bold and italicised values indicate statistically significant comparisons. Sample sizes: Sham = 4 rats, 25 mg/kg = 4 rats, 40 mg/kg = 4 rats, 60 mg/kg = 4 rats. **Abbreviations**: ANOVA, analysis of variance; CI, confidence interval; DFd, degrees of freedom (denominator); DFn, degrees of freedom (numerator); p, probability value.

**Supplementary Table 5.** Results of the statistical analysis for Cleaved Caspase-3 (CC3) cell counts in the infarct border and core.

| **Figure** | **Test name** | **Main effects** | | | **Sidak's multiple comparison** | | | |
|---|---|---|---|---|---|---|---|---|
| | | **Interaction F (DFn, DFd), p value** | **Stroke doses, F (DFn, DFd), p value** | **Variables, F (DFn, DFd), p value** | **Time post-stroke** | **Stroke doses comparisons** | **95% CI** | **p value** |
| **Figure 2A,** CC3 Infarct border | One-way ANOVA | – | F (3, 12) = 4.995, ***p = 0.0178*** | – | 24 h | Sham vs. 25 mg/kg | -126.4 to -10.60 | ***0.0197*** |
| | | | | | | Sham vs. 40 mg/kg | -125.4 to -9.596 | ***0.0215*** |
| | | | | | | Sham vs. 60 mg/kg | -119.2 to -3.346 | ***0.0373*** |

| **Figure** | **Test name** | **Main effects** | | | **Sidak's multiple comparison** | | | |
|---|---|---|---|---|---|---|---|---|
| | | **Interaction F (DFn, DFd), p value** | **Stroke doses, F (DFn, DFd), p value** | **Time post-stroke, F (DFn, DFd), p value** | **Time post-stroke** | **Stroke doses comparisons** | **95% CI** | **p value** |
| **Figure 3A,** CC3 Infarct core | Two-way ANOVA | F (6, 36) = 5.376, ***p = 0.0005*** | F (3, 36) = 16.22, ***p < 0.0001*** | F (2, 36) = 53.94, ***p < 0.0001*** | 24 h | Sham vs. 25 mg/kg | -357.9 to 276.9 | 0.9996 |
| | | | | | | Sham vs. 40 mg/kg | -354.9 to 279.9 | 0.9997 |
| | | | | | | Sham vs. 60 mg/kg | -337.9 to 296.9 | >0.9999 |
| | | | | | | 25mg/kg vs. 40 mg/kg | -314.4 to 320.4 | >0.9999 |
| | | | | | | 25mg/kg vs. 60 mg/kg | -297.4 to 337.4 | >0.9999 |
| | | | | | | 40mg/kg vs. 60 mg/kg | -300.4 to 334.4 | >0.9999 |
| | | | | | 7 d | Sham vs. 25 mg/kg | -650.9 to -16.09 | ***0.0351*** |
| | | | | | | Sham vs. 40 mg/kg | -651.9 to -17.09 | ***0.0343*** |
| | | | | | | Sham vs. 60 mg/kg | -650.7 to -15.84 | ***0.0353*** |
| | | | | | | 25mg/kg vs. 40 mg/kg | -318.4 to 316.4 | >0.9999 |
| | | | | | | 25mg/kg vs. 60 mg/kg | -317.2 to 317.7 | >0.9999 |
| | | | | | | 40mg/kg vs. 60 mg/kg | -316.2 to 318.7 | >0.9999 |

| | | | | | | | | |
|---|---|---|---|---|---|---|---|---|
| | | | | | 14 d | Sham vs. 25 mg/kg | -1211 to -576.1 | ***<0.0001*** |
| | | | | | | Sham vs. 40 mg/kg | -987.9 to -353.1 | ***<0.0001*** |
| | | | | | | Sham vs. 60 mg/kg | -948.2 to -313.3 | ***<0.0001*** |
| | | | | | | 25 mg/kg vs. 40 mg/kg | -94.41 to 540.4 | 0.3026 |
| | | | | | | 25 mg/kg vs. 60 mg/kg | -54.66 to 580.2 | 0.1519 |
| | | | | | | 40 mg/kg vs. 60 mg/kg | -277.7 to 357.2 | 0.9996 |

CC3-positive cell counts in the infarct border and core were analysed using one- or two-way ANOVA, with significance set at $p < 0.05$. Sidak's post hoc test was used for multiple comparisons. Bold and italicised values indicate statistically significant comparisons. Sample sizes: Sham = 4 rats, 25 mg/kg = 4 rats, 40 mg/kg = 4 rats, 60 mg/kg = 4 rats. **Abbreviations:** ANOVA, analysis of variance; CC3, cleaved caspase-3; CI, confidence interval; DFd, degrees of freedom (denominator); DFn, degrees of freedom (numerator); *p*, probability value.

**Supplementary Table 6.** Results of the statistical analysis for Iba1 and GFAP area coverage in the glial reactive zone

| Figure | Test name | Main effects | | | Sidak's multiple comparison | | | |
|---|---|---|---|---|---|---|---|---|
| | | Interaction F (DFn, DFd), p value | Stroke doses, F (DFn, DFd), p value | Variables, F (DFn, DFd), p value | Time post-stroke | Stroke doses comparisons | 95% CI | p value |
| **Figure 2B,** Iba1 Infarct border | One-way ANOVA | – | F (3, 12) = 15.89, ***p = 0.0002*** | – | 24 h | Sham vs. 25 mg/kg | -13.51 to -5.226 | ***0.0001*** |
| | | | | | | Sham vs. 40 mg/kg | -11.84 to -3.553 | ***0.0007*** |
| | | | | | | Sham vs. 60 mg/kg | -11.96 to -3.673 | ***0.0006*** |
| **Figure 2C,** GFAP Infarct border | One-way ANOVA | – | F (3, 12) = 36.89, ***p < 0.0001*** | – | 24 h | Sham vs. 25 mg/kg | -6.705 to -1.290 | ***0.0045*** |
| | | | | | | Sham vs. 40 mg/kg | -9.310 to -3.895 | ***< 0.0001*** |
| | | | | | | Sham vs. 60 mg/kg | -12.64 to -7.227 | ***< 0.0001*** |
| **Supplementary Figure 6,** Iba1 Glial reactive zone width | One-way ANOVA | – | F (2, 9) = 3.085, p = 0.0954 | – | 24 h | 25 mg/kg vs. 40 mg/kg | -28.08 to 70.40 | 0.5622 |
| | | | | | | 25 mg/kg vs. 60 mg/kg | -7.394 to 91.08 | 0.1007 |
| | | | | | | 40 mg/kg vs. 60 mg/kg | -28.56 to 69.92 | 0.5792 |

One-way ANOVA was used to assess the effects of dose on Iba1 and GFAP immunoreactivity in the infarct border and glial reactive zone at 24 h post-injury. Sidak's post hoc test was applied for multiple comparisons. Significance was set at $p < 0.05$. Bold and italicised values indicate statistically significant comparisons**.** Sample sizes: Sham = 4 rats, 25 mg/kg = 4 rats, 40 mg/kg = 4 rats, 60 mg/kg = 4 rats. **Abbreviations:** ANOVA, analysis of variance; CI, confidence interval; DFd, degrees of freedom (denominator); DFn, degrees of freedom (numerator); GFAP, glial fibrillary acidic protein; Iba1, ionised calcium-binding adaptor molecule 1; *p*, probability value.

**Supplementary Table 7.** Results of the statistical analysis for Iba1 and GFAP area coverage in the infarct core

| Figure | Test name | Main effects | | | Sidak's multiple comparison | | | |
|---|---|---|---|---|---|---|---|---|
| | | Interaction F (DFn, DFd), p value | Stroke doses, F (DFn, DFd), p value | Time post-stroke, F (DFn, DFd), p value | Time post-stroke | Stroke doses comparisons | 95% CI | p value |
| **Figure 3B,** Iba1 Infarct core | Two-way ANOVA | F (6, 36) = 10.25, ***p < 0.0001*** | F (3, 36) = 31.78, ***p < 0.0001*** | F (2, 36) = 70.81, ***p < 0.0001*** | 24 h | Sham vs. 25 mg/kg | -11.02 to 10.86 | > 0.9999 |
| | | | | | | Sham vs. 40 mg/kg | -10.87 to 11.00 | > 0.9999 |
| | | | | | | Sham vs. 60 mg/kg | -10.83 to 11.04 | > 0.9999 |
| | | | | | | 25 mg/kg vs. 40 mg/kg | -10.79 to 11.08 | > 0.9999 |
| | | | | | | 25 mg/kg vs. 60 mg/kg | -10.75 to 11.12 | > 0.9999 |
| | | | | | | 40 mg/kg vs. 60 mg/kg | -10.90 to 10.97 | > 0.9999 |
| | | | | | 7 d | Sham vs. 25 mg/kg | -40.59 to -18.72 | ***< 0.0001*** |
| | | | | | | Sham vs. 40 mg/kg | -33.08 to -11.21 | ***< 0.0001*** |
| | | | | | | Sham vs. 60 mg/kg | -33.01 to -11.13 | ***< 0.0001*** |
| | | | | | | 25 mg/kg vs. 40 mg/kg | -3.420 to 18.45 | 0.3264 |
| | | | | | | 25 mg/kg vs. 60 mg/kg | -3.348 to 18.52 | 0.316 |
| | | | | | | 40 mg/kg vs. 60 mg/kg | -10.86 to 11.01 | > 0.9999 |
| | | | | | 14 d | Sham vs. 25 mg/kg | -37.47 to -15.60 | ***< 0.0001*** |
| | | | | | | Sham vs. 40 mg/kg | -30.14 to -8.265 | ***0.0001*** |
| | | | | | | Sham vs. 60 mg/kg | -47.53 to -25.66 | ***< 0.0001*** |
| | | | | | | 25 mg/kg vs. 40 mg/kg | -3.603 to 18.27 | 0.3536 |
| | | | | | | 25 mg/kg vs. 60 mg/kg | -21.00 to 0.8753 | 0.0855 |

| | | | | | | | | |
|---|---|---|---|---|---|---|---|---|
| | | | | | | 40 mg/kg vs. 60 mg/kg | -28.33 to -6.457 | ***0.0005*** |
| **Figure 3C,** GFAP Infarct core | Two-way ANOVA | F (6, 36) = 4.997, ***p = 0.0008*** | F (3, 36) = 19.82, ***p < 0.0001*** | F (2, 36) = 41.89, ***p < 0.0001*** | 24 h | Sham vs. 25 mg/kg | -8.862 to 6.717 | 0.9823 |
| | | | | | | Sham vs. 40 mg/kg | -8.867 to 6.712 | 0.9821 |
| | | | | | | Sham vs. 60 mg/kg | -10.49 to 5.090 | 0.7871 |
| | | | | | | 25 mg/kg vs. 40 mg/kg | -7.795 to 7.785 | > 0.9999 |
| | | | | | | 25 mg/kg vs. 60 mg/kg | -9.417 to 6.162 | 0.9424 |
| | | | | | | 40 mg/kg vs. 60 mg/kg | -9.412 to 6.167 | 0.9429 |
| | | | | | 7 d | Sham vs. 25 mg/kg | -27.65 to -12.07 | ***< 0.0001*** |
| | | | | | | Sham vs. 40 mg/kg | -26.76 to -11.18 | ***< 0.0001*** |
| | | | | | | Sham vs. 60 mg/kg | -26.10 to -10.52 | ***< 0.0001*** |
| | | | | | | 25 mg/kg vs. 40 mg/kg | -6.900 to 8.680 | 0.9867 |
| | | | | | | 25 mg/kg vs. 60 mg/kg | -6.240 to 9.340 | 0.9497 |
| | | | | | | 40 mg/kg vs. 60 mg/kg | -7.130 to 8.450 | 0.9957 |
| | | | | | 14 d | Sham vs. 25 mg/kg | -15.62 to -0.03796 | ***0.0485*** |
| | | | | | | Sham vs. 40 mg/kg | -19.31 to -3.733 | ***0.0017*** |
| | | | | | | Sham vs. 60 mg/kg | -20.34 to -4.760 | ***0.0006*** |
| | | | | | | 25 mg/kg vs. 40 mg/kg | -11.48 to 4.095 | 0.5828 |
| | | | | | | 25 mg/kg vs. 60 mg/kg | -12.51 to 3.067 | 0.3736 |
| | | | | | | 40 mg/kg vs. 60 mg/kg | -8.817 to 6.762 | 0.9844 |

Significance set at p<0.05. The main effects and multiple comparisons using Sidak's post hoc test were reported. Bold + italicised values = statistically significant. Iba1 and GFAP; Rats: Sham controls = 4 pups, 25mg/kg= 4 pups, 40mg/kg= 4 pups, 60mg/kg= 4 pups. **Abbreviations**:

ANOVA, analysis of variance; CI, confidence interval; DFd, degrees of freedom (denominator); DFn, degrees of freedom (numerator); GFAP, glial fibrillary acidic protein; Iba1, ionised calcium-binding adaptor molecule 1; *p*, probability value.

**Supplementary Table 8.** Summary of statistical analysis for behavioural tasks

| Figure | Test name | **Main effects** | | | **Sidak's multiple comparison** | | | |
|---|---|---|---|---|---|---|---|---|
| | | Interaction F (DFn, DFd), p value | Stroke doses, F (DFn, DFd), p value | Variables, F (DFn, DFd), p value | Time post-stroke | Stroke doses comparisons | 95% CI | p value |
| **Figure 4B,** Wire hang | One-way ANOVA | – | F (3, 48) = 3.008, ***p = 0.0393*** | – | 24 h | Sham vs. 25 mg/kg | 0.2746 to 14.05 | ***0.0392*** |
| | | | | | | Sham vs. 40 mg/kg | -0.1718 to 13.24 | 0.0583 |
| | | | | | | Sham vs. 60 mg/kg | -3.694 to 10.08 | 0.5898 |
| **Figure 4C,** Wire hang | One-way ANOVA | – | F (3, 33) = 2.470, *p* = 0.0792 | – | 7 d | Sham vs. 25 mg/kg | -0.6401 to 21.36 | ***0.0699*** |
| | | | | | | Sham vs. 40 mg/kg | -1.650 to 20.35 | 0.1154 |
| | | | | | | Sham vs. 60 mg/kg | -6.005 to 16.00 | 0.5975 |
| **Figure 4D,** Cylinder rearing | Two-way ANOVA | F (6, 51) = 0.4369, *p* = 0.8508 | F (3, 51) = 9.735, ***p < 0.0001*** | F (2, 51) = 1.171, *p* = 0.3182 | 12 d | **Left paw touches** | | |
| | | | | | | Sham vs. 25 mg/kg | 2.098 to 15.90 | ***0.0067*** |
| | | | | | | Sham vs. 40 mg/kg | 1.348 to 15.15 | ***0.0143*** |
| | | | | | | Sham vs. 60 mg/kg | 1.348 to 15.15 | ***0.0143*** |
| | | | | | | **Right paw touches** | | |
| | | | | | | Sham vs. 25 mg/kg | 0.2093 to 14.01 | ***0.0416*** |
| | | | | | | Sham vs. 40 mg/kg | -1.541 to 12.26 | 0.1715 |
| | | | | | | Sham vs. 60 mg/kg | -1.041 to 12.76 | 0.1181 |
| | | | | | | **Both paw touches** | | |
| | | | | | | Sham vs. 25 mg/kg | -1.402 to 12.40 | 0.155 |
| | | | | | | Sham vs. 40 mg/kg | -1.902 to 11.90 | 0.2204 |
| | | | | | | Sham vs. 60 mg/kg | -4.402 to 9.402 | 0.7563 |
| **Figure 4E,** Cylinder rearing | One-way ANOVA | – | F (3, 17) = 5.190, ***p = 0.0100*** | – | 12 d | Sham vs. 25 mg/kg | 2.356 to 26.09 | ***0.0166*** |
| | | | | | | Sham vs. 40 mg/kg | 1.441 to 25.17 | ***0.0256*** |
| | | | | | | Sham vs. 60 mg/kg | -0.9988 to 22.73 | 0.0782 |

| | | | | | | | | |
|---|---|---|---|---|---|---|---|---|
| **Figure 4F,** Adhesive tape removal | One-way ANOVA | – | F (3, 17) = 2.947, p=0.0625 | – | 12 d | Sham vs. 25 mg/kg | -79.76 to -2.450 | ***0.0354*** |
| | | | | | | Sham vs. 40 mg/kg | -57.84 to 19.47 | 0.5002 |
| | | | | | | Sham vs. 60 mg/kg | -64.13 to 13.18 | 0.2689 |
| **Figure 4G,** Adhesive tape removal | One-way ANOVA | – | F (3, 17) = 8.183, ***p = 0.0014*** | – | 12 d | Sham vs. 25 mg/kg | -1.260 to -0.2687 | ***0.0023*** |
| | | | | | | Sham vs. 40 mg/kg | -0.9571 to 0.033 | 0.0719 |
| | | | | | | Sham vs. 60 mg/kg | -1.217 to -0.2262 | ***0.0038*** |

Behavioural data were analysed using one-way or two-way ANOVA, with significance set at p<0.05. The main effects and multiple comparisons using Sidak's post hoc test were reported. Bold + italicised values = statistically significant. 24h wire hang; Rats: Sham controls = 18 pups, 25mg/kg= 11 pups, 40mg/kg= 12 pups, 60mg/kg= 11 pups. 7d wire hang; Rats: Sham controls = 13 pups, 25mg/kg= 8 pups, 40mg/kg= 8 pups, 60mg/kg= 8 pups. Cylinder rearing and adhesive tape removal; Rats: Sham controls = 9 pups, 25mg/kg= 4 pups, 40mg/kg= 4 pups, 60mg/kg= 4 pups.

**Supplementary Table 9.** Exploratory correlations between behavioural outcomes and lesion volumes.

| | **Pearson r** | **$R^2$** | ***p* value** |
|---|---|---|---|
| **Wire Hang Time at 24 hours vs. lesion volume at 24 hours** | | | |
| **25 mg/kg RB** | –0.526 | 0.277 | 0.236 |
| **40 mg/kg RB** | –0.718 | 0.515 | 0.245 |
| **60 mg/kg RB** | 0.029 | 0.0008 | 0.490 |
| **25 and 40 mg/kg** | –0.633 | 0.400 | *0.063* |
| **25 and 40 and 60 mg/kg** | 0.330 | 0.1091 | 0.175 |
| **Wire Hang Time at 24 hours vs lesion volume at 7 days** | | | |
| **25 mg/kg RB** | –0.813 | 0.661 | *0.093* |
| **40 mg/kg RB** | –0.995 | 0.991 | *0.002* |
| **60 mg/kg RB** | 0.326 | 0.106 | 0.336 |
| **12-day contralateral/ipsilateral paw use versus lesion volume at 14 days** | | | |
| **25 mg/kg RB** | 0.946 | 0.895 | *0.026* |
| **40 mg/kg RB** | –0.167 | 0.028 | 0.416 |
| **60 mg/kg RB** | –0.161 | 0.026 | 0.419 |

**Abbreviations:** RB: Rose Bengal

**Supplementary Table 10.** Summary of sex-specific statistical analysis for the 25 mg/kg stroke dose

This table presents the results of two-way ANOVA analyses assessing the main and interaction effects of sex, injury, and time post-stroke on histological (infarct volume, cleaved caspase-3, GFAP, Iba1) and behavioural (wire hang) outcomes following a 25 mg/kg Rose Bengal–induced focal ischemic injury. Sidak's multiple comparisons were applied to explore sex- and injury-related differences across time points (24 h, 7 d, 14 d). Reported values include F(DFn, DFd), corresponding p-values, 95% confidence intervals (CI), and post hoc pairwise comparisons.

| ***Figure*** | ***Test name*** | **Main effects** | | | **Sidak's multiple comparison** | | | |
|---|---|---|---|---|---|---|---|---|
| | | ***Interaction F (DFn, DFd), p value*** | ***Sex, F (DFn, DFd), p value*** | ***Time post-stroke, F (DFn, DFd), p value*** | ***Time post-stroke*** | ***Sex/injury comparisons*** | ***95% CI*** | ***p value*** |
| **Figure 5A,** Infarct volume | Two-way ANOVA | F (2, 18) = 0.9857, p = 0.3924 | F (1, 18) = 1.034, p = 0.3227 | F (2, 18) = 38.46, ***px < 0.0001*** | 24h | Males - Females | -0.563 to 2.696 | 0.2766 |
| | | | | | 7d | | -1.529 to 1.731 | 0.9979 |
| | | | | | 14d | | -1.706 to 1.554 | 0.9991 |
| **Figure** | **Test name** | **Interaction F (DFn, DFd), p value** | **Sex, F (DFn, DFd), p value** | **Injury, F (DFn, DFd), p value** | **Injury/Sex** | **Sex/injury comparisons** | **95% CI** | **p value** |
| **Figure 5B,** Wire hang 24h | Two-way ANOVA | F (1, 39) = 0.1337, p = 0.7166 | F (1, 39) = 0.6780, p = 0.4153 | F (1, 39) = 14.54, ***p = 0.0005*** | Sham | Males - Females | -3.831 to 8.564 | 0.6155 |
| | | | | | Stroke | | -5.967 to 7.789 | 0.9422 |
| | | | | | Males | Sham - Stroke | 1.623 to 15.01 | ***0.0125*** |
| | | | | | Fem. | | 0.4661 to 13.26 | ***0.0336*** |
| **Figure 5B,** Wire hang 7d | Two-way ANOVA | F (1, 32) = 0.3341, *p* = 0.5673 | F (1, 32) = 0.02398, *p* = 0.8779 | F (1, 32) = 4.619, ***p = 0.0393*** | Sham | Males - Females | -11.65 to 8.809 | 0.9357 |
| | | | | | Stroke | | -9.545 to 14.47 | 0.8657 |
| | | | | | Males | Sham - Stroke | -6.411 to 16.98 | 0.5060 |
| | | | | | Fem. | | -1.422 to 19.76 | 0.0986 |
| **Figure 5C,** CC3 Infarct border | Two-way ANOVA | F (1, 12) = 0.03197, *p* = 0.8611 | F (1, 12) = 0.001812, *p* = 0.9667 | F (1, 12) = 14.22, ***p = 0.0027*** | Sham | Males - Females | -56.26 to 49.76 | 0.9852 |
| | | | | | Stroke | | -51.01 to 55.01 | 0.9944 |
| | | | | | Males | Sham - Stroke | -111 to -4.988 | ***0.0322*** |

| | | | | | | | | |
|---|---|---|---|---|---|---|---|---|
| 24h | | | | | Fem. | | -105.8 to 0.262 | 0.0512 |
| **Figure 5D,** Iba1 Infarct border 24h | Two-way ANOVA | F (1, 12) = 5.866e, *p* = 0.9940 | F (1, 12) = 0.01790, *p* = 0.8958 | F (1, 12) = 148.4, *p* < 0.0001 | Sham | Males - Females | -3.123 to 2.888 | 0.9939 |
| | | | | | Stroke | | -3.110 to 2.900 | 0.9952 |
| | | | | | Males | Sham - Stroke | -13.15 to -7.14 | ***< 0.0001*** |
| | | | | | Fem. | | -13.14 to -7.12 | ***< 0.0001*** |
| **Figure 5E,** GFAP Infarct border 24h | Two-way ANOVA | F (1, 12) = 0.1033, *p* = 0.7535 | F (1, 12) = 0.1303, *p* = 0.7243 | F (1, 12) = 221.5, ***p* < 0.0001** | Sham | Males - Females | -1.125 to 1.150 | 0.9995 |
| | | | | | Stroke | | -0.9226 to 1.35 | 0.8690 |
| | | | | | Males | Sham - Stroke | -5.928 to -3.65 | ***< 0.0001*** |
| | | | | | Fem. | | -5.725 to -3.45 | ***< 0.0001*** |
| **Figure** | **Test name** | **Interaction F (DFn, DFd), *p* value** | **Injury/Sex, F (DFn, DFd), *p* value** | **Time post-stroke, F (DFn, DFd), *p* value** | **Time post-stroke** | **Sex/injury comparisons** | **95% CI** | **p value** |
| **Figure 5D,** Ib1 Infarct core | Two-way ANOVA | F (6, 36) = 33.66, ***p* < 0.0001** | F (3, 36) = 131.1, ***p* < 0.0001** | F (2, 36) = 103.8, ***p* < 0.0001** | 24 h | Sham Males vs. Stroke Males | -6.861 to 7.406 | > 0.9999 |
| | | | | | | Sham Males vs. Sham Females | -7.251 to 7.016 | > 0.9999 |
| | | | | | | Sham Males vs. Stroke Females | -7.479 to 6.789 | > 0.9999 |
| | | | | | | Stroke Males vs. Sham Females | -7.524 to 6.743 | > 0.9999 |
| | | | | | | Stroke Males vs. Stroke Females | -7.751 to 6.516 | > 0.9999 |
| | | | | | | Sham Females vs. Stroke Females | -7.361 to 6.906 | > 0.9999 |
| | | | | | 7 d | Sham Males vs. Stroke Males | -35.41 to -21.1 | ***< 0.0001*** |
| | | | | | | Sham Males vs. Sham Females | -7.651 to 6.616 | > 0.9999 |
| | | | | | | Sham Males vs. Stroke Females | -42.42 to -28.1 | ***< 0.0001*** |
| | | | | | | Stroke Males vs. Sham | 20.62 to 34.89 | ***< 0.0001*** |

| | | | | | | | | |
|---|---|---|---|---|---|---|---|---|
| | | | | | | Females | | |
| | | | | | | Stroke Males vs. Stroke Females | -14.14 to 0.123 | 0.0563 |
| | | | | | | Sham Females vs. Stroke Females | -41.90 to -27.6 | ***< 0.0001*** |
| | | | | | 14 d | Sham Males vs. Stroke Males | -38.16 to -23.8 | ***< 0.0001*** |
| | | | | | | Sham Males vs. Sham Females | -6.741 to 7.526 | > 0.9999 |
| | | | | | | Sham Males vs. Stroke Females | -36.88 to -22.6 | ***< 0.0001*** |
| | | | | | | Stroke Males vs. Sham Females | 24.28 to 38.55 | ***< 0.0001*** |
| | | | | | | Stroke Males vs. Stroke Females | -5.854 to 8.414 | 0.997 |
| | | | | | | Sham Females vs. Stroke Females | -37.27 to -23.0 | ***< 0.0001*** |
| **Figure 5E,** GFAP Infarct core | Two-way ANOVA | F (6, 36) = 15.42, ***p < 0.0001*** | F (3, 36) = 56.22, ***p < 0.0001*** | F (2, 36) = 50.11, ***p < 0.0001*** | 24 h | Sham Males vs. Stroke Males | -6.783 to 4.543 | 0.9949 |
| | | | | | | Sham Males vs. Sham Females | -5.651 to 5.676 | > 0.9999 |
| | | | | | | Sham Males vs. Stroke Females | -6.543 to 4.783 | 0.9987 |
| | | | | | | Stroke Males vs. Sham Females | -4.531 to 6.796 | 0.9946 |
| | | | | | | Stroke Males vs. Stroke Females | -5.423 to 5.903 | > 0.9999 |
| | | | | | | Sham Females vs. Stroke Females | -6.556 to 4.771 | 0.9985 |
| | | | | | 7 d | Sham Males vs. Stroke Males | -26.18 to -14.8 | ***< 0.0001*** |
| | | | | | | Sham Males vs. Sham Females | -5.634 to 5.692 | >0.9999 |
| | | | | | | Sham Males vs. Stroke | -26.20 to -14.8 | ***< 0.0001*** |

| | | | | | | | | |
|---|---|---|---|---|---|---|---|---|
| | | | | | | Females | | |
| | | | | | | Stroke Males vs. Sham Females | 14.88 to 26.21 | ***< 0.0001*** |
| | | | | | | Stroke Males vs. Stroke Females | -5.681 to 5.646 | > 0.9999 |
| | | | | | | Sham Females vs. Stroke Females | -26.23 to -14.9 | ***< 0.0001*** |
| | | | | | 14 d | Sham Males vs. Stroke Males | -16.30 to -4.96 | ***< 0.0001*** |
| | | | | | | Sham Males vs. Sham Females | -6.358 to 4.968 | 0.9997 |
| | | | | | | Sham Males vs. Stroke Females | -17.33 to -6.00 | ***< 0.0001*** |
| | | | | | | Stroke Males vs. Sham Females | 4.274 to 15.60 | ***0.0001*** |
| | | | | | | Stroke Males vs. Stroke Females | -6.701 to 4.626 | 0.9967 |
| | | | | | | Sham Females vs. Stroke Females | -16.64 to -5.31 | ***< 0.0001*** |
| **Figure 5C,** CC3 Infarct core | Two-way ANOVA | F (6, 36) = 26.47, ***p < 0.0001*** | F (3, 36) = 65.83, ***p < 0.0001*** | F (2, 36) = 90.40, ***p < 0.0001*** | 24 h | Sham Males vs. Stroke Males | -226.8 to 188.8 | > 0.9999 |
| | | | | | | Sham Males vs. Sham Females | -211.0 to 204.5 | > 0.9999 |
| | | | | | | Sham Males vs. Stroke Females | -217 to 198.5 | > 0.9999 |
| | | | | | | Stroke Males vs. Sham Females | -192 to 223.5 | > 0.9999 |
| | | | | | | Stroke Males vs. Stroke Females | -198 to 217.5 | > 0.9999 |
| | | | | | | Sham Females vs. Stroke Females | -213.8 to 201.8 | > 0.9999 |
| | | | | | 7 d | Sham Males vs. Stroke Males | -595.8 to -180. | ***< 0.0001*** |
| | | | | | | Sham Males vs. Sham | -213.5 to 202 | > 0.9999 |

| | | | | | | | | |
|---|---|---|---|---|---|---|---|---|
| | | | | | | Females | | |
| | | | | | | Sham Males vs. Stroke Females | -487.5 to -71.9 | ***0.0037*** |
| | | | | | | Stroke Males vs. Sham Females | 174.5 to 590 | **< *0.0001*** |
| | | | | | | Stroke Males vs. Stroke Females | -99.55 to 316 | 0.6378 |
| | | | | | | Sham Females vs. Stroke Females | -481.8 to -66.2 | ***0.0047*** |
| | | | | | 14 d | Sham Males vs. Stroke Males | -1155 to -739.7 | **< *0.0001*** |
| | | | | | | Sham Males vs. Sham Females | -178.5 to 237 | 0.9992 |
| | | | | | | Sham Males vs. Stroke Females | -1102 to -686.7 | **< *0.0001*** |
| | | | | | | Stroke Males vs. Sham Females | 769 to 1185 | **< *0.0001*** |
| | | | | | | Stroke Males vs. Stroke Females | -154.8 to 260.8 | 0.9807 |
| | | | | | | Sham Females vs. Stroke Females | -1132 to -716 | **< *0.0001*** |

Significance set at $p < 0.05$. The main effects and multiple comparisons using Sidak's post hoc test were reported. Bold + italicised values = statistically significant. Infarct volume, Iba1 and GFAP area coverage (Figure 8A, 8D–G): 4 rat pups per group. 24h wire hang; Rats: Sham controls- Males = 10 pups, Females = 14 pups; Stroke - Males = 10 pups, Females = 9 pups. 7d wire hang; Rats: Sham controls - Males = 9 pups, Females = 12 pups; Stroke- Males = 7 pups, Females = 8 pups. **Abbreviations:** CC3, cleaved caspase-3; CI, confidence interval; DFd, degrees of freedom (denominator); DFn, degrees of freedom (numerator); GFAP, glial fibrillary acidic protein; Iba1, ionised calcium-binding adaptor molecule 1; P, postnatal day; *p*, probability value.

**Table 11.** Power calculation and effect size for lesion volume and behavioural outcomes by sex.

| | **Males Mean ± SD** | **Females Mean ± SD** | **Cohen's d** | **Interpretation** | ***N* per group for p < 0.05** |
|---|---|---|---|---|---|
| ***Lesion volume analyses over time (male vs female)*** | | | | | |
| **24 h** | 4.35 ± 1.85 | 3.28 ± 0.41 | 0.75 | Large | 29 |
| **7 d** | 0.93 ± 0.66 | 0.83 ± 0.29 | 0.22 | Small | 385 |
| **14 d** | 0.16 ± 0.13 | 0.24 ± 0.18 | 0.44 | Medium | 80 |
| ***Wire hang at 24 hours (sham vs injured by sex)*** | | | | | |
| **Sham** | 17.05 ± 7.39 | 15.31 ± 7.96 | 0.23 | Small | 297 |
| **Stroke** | 8.43 ± 2.71 | 7.70 ± 2.85 | 0.26 | Small | 232 |
| ***Wire hang at 7 days (sham vs injured by sex)*** | | | | | |
| **Sham** | 18.58 ± 9.28 | 20.20 ± 13.11 | 0.14 | Small | 801 |
| **Stroke** | 14.77 ± 8.58 | 12.74 ± 3.99 | 0.30 | Small | 174 |

**Abbreviations**: *d* (Cohen's d), effect size; *d,* days; *h*, hours; *N*, total sample size; SD, standard deviation; *vs*, versus.
